# Phylogeny-Guided Global–Local Shrinkage for Microbiome Feature Selection

**DOI:** 10.64898/2026.10.28.762580

**Authors:** Zihan Zhu, Shuangjie Zhang, Xueying Tang, Liangliang Zhang

## Abstract

Microbiome feature selection is challenging because sequencing counts are sparse and compositional, while evolutionarily related taxa may exhibit similar associations with an exposure. We propose phylogeny-guided global–local shrinkage (PGL), a Bayesian regression method for identifying exposure-associated differences in the relative abundances of individual amplicon sequence variants (ASVs). A multinomial logit-normal working model analyzes relative abundance among ASVs detected in each sample. A zero-sum constraint gives the regression coefficients a log-ratio interpretation relative to the mean across retained ASVs. Global–local shrinkage accommodates weak and large associations, while an estimated weight controls information sharing along a supplied phylogenetic tree. Simulation ablation studies showed that phylogenetic borrowing improved detection power and estimation of nonzero coefficients when associations aligned with the tree, with little change under a tree-independent generator. Estimated borrowing decreased as the supplied tree was corrupted. Comparisons with benchmark methods revealed different trade-offs between detection power and false discovery rates across simulation settings. In an HIV oral microbiome application, ablation analysis showed that phylogenetic borrowing preserved the overall pattern of estimated associations while changing posterior uncertainty and selecting three additional ASVs. PGL provides ASV-level inference with adaptive information sharing across evolutionarily related features.

## 1 Introduction

Microbiome studies often seek to identify microbial features associated with a host phenotype or exposure. In 16S rRNA data, amplicon sequence variants (ASVs) retain sequence-level resolution and can be paired with an inferred phylogenetic tree (Callahan et al., 2016; Callahan, McMurdie, and Holmes, 2017). Aggregating ASVs before analysis may conceal different responses within a named taxon, whereas treating all ASVs as unrelated ignores the possibility that evolutionary neighbors respond similarly. The statistical objective is to retain ASV-specific inference while allowing related features to inform one another.

Phylogeny provides a biologically motivated source of information, but its relevance may vary. Microbial traits differ in their degree of phylogenetic conservation, so related organisms may share some characteristics without exhibiting similar responses to an exposure (Martiny et al., 2015). This distinction matters when a tree is used to guide statistical inference, since even closely related ASVs may have associations that differ in magnitude or direction. A prior that borrows information without imposing equality offers an alternative, preserving ASV-specific effects while allowing shared structure to inform their estimation. The strength of this borrowing should be learned from the data rather than fixed in advance, because the usefulness of a supplied tree may vary across exposures and datasets.

The count table creates additional challenges. Sequencing counts within a sample share a finite sequencing depth, and sequencing depth can vary across samples. Consequently, changes in relative abundance cannot generally be interpreted as changes in absolute abundance without additional information (Aitchison, 1982; McMurdie and Holmes, 2014). Feature selection therefore requires a defined compositional reference: a nonzero coefficient represents a relative association, and changing that reference can change which features are selected. The reference must be specified to interpret an ASV’s selection and the direction of its association.

Zeros are common in the microbiome count table and complicate both modeling and interpretation. A zero count can reflect biological absence, limited sequencing depth, or technical variation, and the count alone does not identify its cause. Replacing zeros by a pseudo-count permits log transformations, but changes the values entering an abundance comparison. An alternative is to analyze relative abundance on the observed support. This approach avoids replacing zero counts and targets abundance associations with support treated as fixed; occurrence associations and the causes of non-detection remain separate questions.

Several established approaches address aspects of these problems. DESeq2 uses negative binomial regression with normalization, while LinDA and ALDEx2 address compositional associations through log-ratio analyses (Fernandes et al., 2014; Love, Huber, and Anders, 2014; Zhou et al., 2022). ANCOM-BC2 supports covariate-adjusted inference across groups, and MaAsLin 3 distinguishes prevalence from nonzero-abundance associations (Lin and Peddada, 2024; Nickols et al., 2026). Other approaches combine normalization with zero-inflated modeling (Paulson et al., 2013) or introduce variable selection in Dirichlet–multinomial regression (Chen and Li, 2013). These methods differ in their treatment of zeros, reference construction, and inferential targets.

Related work uses microbial features as predictors of an outcome. In Bayesian compositional regression, Zhang et al. (2021) combine a zero-constrained coefficient prior with an Ising prior that encourages joint selection of features related by genetic sequence similarity. For sparse predictors more generally, Yan and Bien (2021) develop tree-guided aggregation of rare features to improve prediction. Building on this approach, Bien et al. (2021) introduce tree-aggregated compositional regression (trac), which learns aggregation levels within a log-contrast predictive model. These approaches use structure to inform predictor selection and aggregation. In the present study, microbial counts form the multivariate response, and the coefficients describe their associations with exposures.

Other related work uses multivariate models or hierarchical structure to share information across microbial features. Bayesian models have incorporated sparse dependence among count features, while phylogenetically informed graphical models use evolutionary relationships in the estimation of microbial association networks (Chung, Gaynanova, and Ni, 2022; Zhang et al., 2023). In differential abundance testing, StructFDR uses tree-derived relationships to share information among feature-level hypotheses, whereas treeclimbR selects a data-dependent resolution among hypotheses associated with leaves and branches (Huang et al., 2021; Xiao, Cao, and Chen, 2017). Together, these approaches show that structure can inform dependence estimation, variable selection, aggregation, multiple testing, and the resolution of reported findings. Our focus is adaptive joint shrinkage of regression effects along a supplied tree, with inference still reported for individual ASVs under an explicitly conditional abundance estimand.

The methodological challenge is to combine shrinkage, phylogenetic borrowing, and a clearly identified compositional target. Shrinkage is useful when many associations are weak and individual ASVs have limited information, but excessive shrinkage can obscure large effects. A single scale for all features cannot readily accommodate both strongly shrunk and large effects. Global–local priors address this through an overall shrinkage scale and feature-specific scales that permit departures from it. Heavy-tailed scale priors provide substantial flexibility for these departures (Zhu and Tang, 2025; Zhu, Tang, and Zhou, 2026). Introducing phylogenetic dependence requires retaining feature-specific flexibility while preserving a defined compositional reference. The dependence structure, local scales, and centering constraint must therefore be coordinated, with the strength of borrowing allowed to decrease when the tree provides little information about exposure effects.

We develop an observed-support multinomial logit-normal working model with phylogeny-guided global–local shrinkage (PGL). The multinomial component uses the detected ASVs as its support, and a Gaussian regression layer relates latent log weights to exposures and adjustment covariates. An exact zero-sum constraint sets the reference for each exposure coefficient as a model-based log-ratio slope relative to the mean effect across retained ASVs, with support treated as fixed. A global scale encourages shrinkage toward zero, and local scales allow individual coordinates to depart from that shrinkage before centering. A correlation matrix derived from a Brownian model on the supplied tree links standardized latent effects before scaling and centering. An estimated weight controls the strength of this phylogenetic correlation. Thus the tree guides regularization without requiring neighboring ASVs to have identical coefficients. Fixing the weight at zero gives the nested phylogeny-independent model (PIND), which retains the same likelihood, scale priors, and compositional constraint. Comparing PGL with PIND isolates the contribution of phylogenetic borrowing within this framework.

We develop posterior computation and evaluate the method in three complementary simulation experiments. Effects aligned with the tree assess the benefit of informative structure; a separate tree-independent generator examines performance under an external count model and permits comparison with feature-wise methods; and a corrupted input tree tests whether estimated borrowing responds to misspecification. We then apply the method to salivary microbiome data from the HIV study of Jayaraman et al. (2025), examining how phylogenetic borrowing changes ASV-level effect estimates, uncertainty, and selection after covariate adjustment. Sections 2–5 present the model, simulations, application, and discussion, respectively.

## 2 Model

### 2.1 Observed-Support Model and Effect Interpretation

For sample *i* = 1, … , *n*, let ***y***_*i*_ = (*y*_*i*1_, … , *y*_*iK*_)^⊤^ be the counts for *K* amplicon sequence variants (ASVs) and *N*_*i*_ = ∑_*k*_ *y*_*ik*_ the sequencing depth. Let *u*_*ik*_ denote a latent log weight. We condition on the observed support *η*_*ik*_ = *I*(*y*_*ik*_ *>* 0) and use the gated multinomial factor

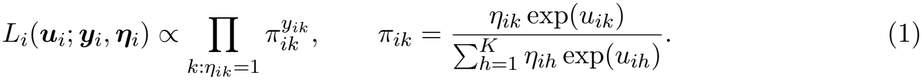

Only ASVs observed in sample *i* enter its compositional denominator. The latent log weights follow

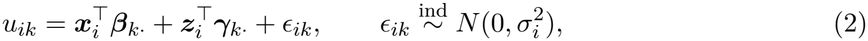

where ***x***_*i*_ contains the exposures of interest and ***z***_*i*_ contains nuisance covariates, including an intercept when appropriate. For ASV *k, β*_*kj*_ and *γ*_*kl*_ are the coefficients of exposure column *j* and nuisance column *l*, respectively, and 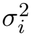 is the residual variance shared by the latent coordinates in sample *i*. The Gaussian layer is defined for all *K* latent coordinates; *η*_*ik*_ gates only the multinomial likelihood. Equations (1) and (2) form an observed-support multinomial logit-normal working model (Aitchison and Shen, 1980).

Because ***η***_*i*_ is computed from the counts and then held fixed, this factor is a working likelihood on the observed support. It does not include the normalizing factor for a multinomial distribution truncated to positive counts in every active category (Appendix C). It does not model presence or distinguish biological absence from a sampling zero. The exposure effects below concern relative abundance in this working model, rather than prevalence or marginal counts. We discuss a joint prevalence–abundance extension in Section 5.

For any two ASVs observed in the same sample, log(*π*_*ik*_*/π*_*ih*_) = *u*_*ik*_ *− u*_*ih*_. Within the working model with support treated as fixed, the expected log ratio has slope *β*_*kj*_ *−β*_*hj*_ in exposure *j*, holding other covariates fixed. The multinomial likelihood is invariant to a common shift of all active *u*_*ik*_, so an unconstrained coefficient level would be anchored by its prior rather than by these ratios. We therefore impose 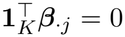 for each exposure column *j*. Our exact constraint chooses the across-ASV mean as the reference: *β*_*kj*_ = *K*^−1^ ∑_*h*_(*β*_*kj*_ *− β*_*hj*_). Thus *β*_*kj*_ is a model-based log-ratio slope relative to the mean effect over the *K* retained ASVs, not an absolute-abundance effect. Changing the ASV filtering rule changes this reference and hence the feature-selection target. Interpretation requires adequate co-observation and covariate information; the prior supplies regularization where those data are weak. Appendix C gives the identification argument.

### 2.2 Adaptive Phylogenetic Correlation

The tree enters through the correlation of a standardized latent vector 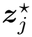 before scaling and projection; the complete prior is specified in Section 2.3.

Let *T* be the supplied rooted phylogeny, with tips matched to the *K* ASVs. Under a Brownian-motion model on *T* , the covariance between two tips is the length of their shared path from the root to their most recent common ancestor. This shared-path covariance arises from independent increments accumulated along branches. We standardize the covariance returned by `ape::vcv.phylo` to a correlation matrix ***R***_phylo_ (Paradis and Schliep, 2019); numerical stabilization preserves unit diagonal and positive definiteness (Appendix A).

An exposure group *g* indexes columns that share a borrowing weight *ω*_*g*_. For a categorical exposure, all non-reference indicators enter the same linear predictor, with separate ***β***_*·j*_ and scales but a shared *ω*_*g*_. Contrasts between its levels are formed from the corresponding coefficient draws. Continuous exposures enter as ordinary columns of ***x***_*i*_. For exposure group *g*, define

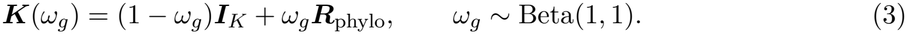

For exposure column *j* in group *g*(*j*), take 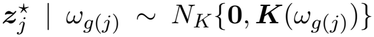. The unit diagonal makes *ω*_*g*_ control phylogenetic correlation among the pre-projection 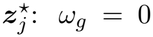 removes that correlation, whereas *ω*_*g*_ = 1 uses the full Brownian correlation. Intermediate values allow borrowing to vary with the data; when information about the weight is limited, its posterior can remain broad. Brownian shared-path correlations are nonnegative: the tree encodes similarity, while the compositional contrast can induce either sign of dependence among the final effects.

### 2.3 Projected Global–Local Prior and Inference

Define **Λ**_*j*_ = diag_*k*_(*λ*_*kj*_) and 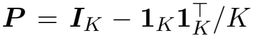. Let 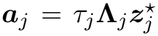 be an unconstrained latent effect vector. The coefficients entering Equation (2) are

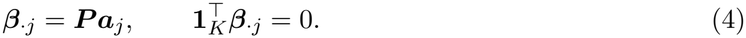

This construction induces a prior on the zero-sum contrast space. Because *β*_*kj*_ *− β*_*hj*_ = *a*_*kj*_ *− a*_*hj*_, projection preserves every pairwise log-ratio slope while removing the common shift.

The global scale *τ*_*j*_ shrinks an exposure’s effect vector, while the local scales *λ*_*kj*_ allow individual latent coordinates to depart from that shrinkage. To accommodate small scales and large latent departures, we use independent positive log-Cauchy priors for both:

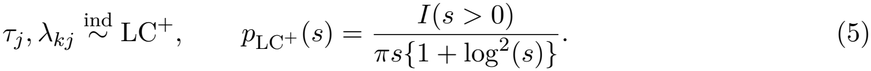

Equivalently, log *s ∼* Cauchy(0, 1). The density has an integrable pole at zero and a very heavy upper tail; the global scale supports shrinkage, while the local scales allow related ASVs to retain different coefficients (Zhu and Tang, 2025; Zhu et al., 2026). The prior does not create exact zeros or taxonomic aggregates.

Conditional on the scales and *ω*_*g*(*j*)_, the final coefficient vector has a Gaussian distribution on the (*K −* 1)-dimensional zero-sum contrast space. Its conditional covariance is

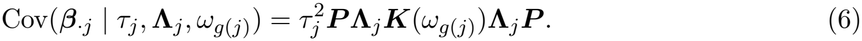

The phylogeny-independent comparator (PIND) fixes every *ω*_*g*_ = 0 and otherwise retains the same model. Even at *ω*_*g*_ = 0, projection creates dependence among the *β*_*kj*_; PIND means independent *pre-projection* latent effects, not independent final contrasts. A change to one local scale affects every centered coordinate. Specifically, changing *a*_*kj*_ by Δ while holding the other coordinates fixed changes *β*_*kj*_ by (1 − 1*/K*)Δ and each *β*_*hj*_, *h ≠ k*, by −Δ*/K*. Thus, the local scales act before centering and do not control the final coefficients in isolation. Phylogenetic borrowing acts through ***K***(*ω*_*g*_) before centering and need not make the final covariances all positive. A large *ω*_*g*_ need not make neighboring final coefficients similar because local scales vary and projection changes their dependence. Appendix B gives the derivation.

The independent nuisance priors are

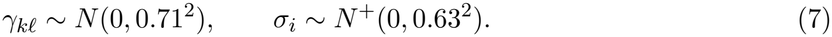

Their calibration is given in Appendix D.

We select ASV *k* for exposure *j* when its 95% equal-tailed posterior credible interval for the identifiable contrast *β*_*kj*_ excludes zero. This rule asks whether the ASV differs from the across-ASV reference; it does not test for a prevalence change or for an absolute abundance change. It does not provide a formal false-discovery-rate guarantee. Posterior computation uses a Metropolis-within-Gibbs sampler with a blocked Gaussian update for 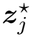 and log-scale updates for the positive scales. Each update preserves ***β***_*·j*_ = ***P a***_*j*_. The full conditionals and acceptance ratios are in Appendix E.

## 3 Simulation

### 3.1 Design and evaluation

Three experiments evaluated the benefit of phylogenetic borrowing when effects follow the tree (Experiment 1), performance against feature-wise benchmarks under a different, tree-independent data generator (Experiment 2), and adaptation to an incorrect input tree (Experiment 3). Every setting used *K* = 200 ASVs and 100 completed Monte Carlo replicates. PGL and PIND shared the likelihood, identifiability constraint, shrinkage priors, and sampler; PIND fixed *ω* = 0. DESeq2 (Love et al., 2014) and the composition-aware LinDA method (Zhou et al., 2022) supplied feature-wise comparisons in Experiments 1 and 2.

Experiment 1 was model-based: it generated counts from a gated multinomial logit-normal mechanism on a random tree. A Gaussian-probit presence process thresholded correlated Gaussian variables to set marginal presence probability *P ∈ {*0.25, 0.50*}*. Among 200 exposure effects, exactly 60 were nonzero; the active effects summed to zero and had root mean square (RMS) 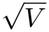, with *V ∈ {*0.5, 1*}*. Alignment *A ∈ {*0, 0.5, 1*}* assigned 0, 30, or 60 active ASVs to coherent clades, leaving the rest randomly located. We crossed these factors at *n* = 50 (12 settings) and added *n* = 100, *P* = 0.5, *V* = 1 at *A* = 0, 1 as a targeted sample-size check. The active-set size and RMS were held fixed across alignment levels. The generator used a correlation derived from the normalized graph-Laplacian pseudoinverse for presence and within-clade effect deviations; the fitted prior used Brownian shared-path correlations from the same tree (Appendix F). Each sample had 50,000 reads, and the generated presence indicators were supplied to both Bayesian fits.

Experiment 2 used a separate LinDA-inspired log-normal absolute-abundance generator followed by multinomial sequencing. It changed the abundance and zero-generating mechanisms as well as removing tree alignment, so its comparison with Experiment 1 does not isolate any one of those changes. The binary exposure depended on two adjusted covariates; a heavy-tailed baseline abundance curve and negative-binomial library sizes produced sampling zeros. The set of ASVs with nonzero coefficients and their coefficient values were generated independently of a random tree, which was supplied only to PGL. Active proportions *S ∈ {*0.10, 0.30*}* were crossed with *V ∈ {*0.5, 1*}* at *n* = 50, and with *V* = 1 at *n* = 100, 200 (eight settings). All taxa were structurally present, and the Bayesian fits used *η*_*ik*_ = 1, including for ASVs with observed sampling zeros.

Experiment 3 held the generated data and true effects fixed within each replicate while degrading only the tree supplied to the fit. It used the fully aligned Experiment 1 generator at *n* = 50, *V* = 1, and both presence regimes. For *q ∈ {*0, 0.25, 0.5, 1*}*, we selected a fraction *q* of the tip labels and permuted them in the fitted Brownian correlation, keeping the topology and branch lengths fixed. The realized correlation between the unpermuted and supplied Brownian kernels averaged approximately 1.00, 0.61, 0.30, 0.00 across these four levels. At every *q* we fitted adaptive PGL with estimated *ω* and a forced-borrowing version with *ω* = 1; PIND was fitted once per replicate as their common baseline. Thus the design separates the benefit of a correct tree from the cost of fixing the borrowing weight at one when the tree is misspecified.

Bayesian fits used 40,000 Markov chain Monte Carlo (MCMC) iterations, discarded the first 20,000 as burn-in, and retained every fourth remaining draw, leaving 5,000 draws. Primary selection required a 95% equal-tailed credible interval for the identifiable *β*_*k*_ to exclude zero; benchmark selection used Benjamini–Hochberg adjusted *p <* 0.05. The two selection rules are not calibrated to the same empirical FDR, so benchmark comparisons describe the observed sensitivity–error trade-off rather than a common error-controlled operating point. We evaluated power, false-positive rate (FPR), false discovery rate (FDR), *F*_1_, and root mean squared error (RMSE) against the zero-sum truth. All selection metrics used the same true active set (Appendix F). FDR is the mean of replicate-level false discovery proportions (FDPs), with empty selections assigned zero. Paired RMSE ratios compare models on the same replicate; values below one favor the numerator. Figures show approximate 95% Monte Carlo intervals for means. Complete designs, all settings, and available sampler diagnostics are in Appendix F.

### 3.2 Experiment 1: aligned signals and sparse support

Phylogenetic borrowing helped when the active effects clustered on the tree, and its benefit increased with alignment (Figure 1, Panels A–B). PGL and PIND had nearly the same power and RMSE over ASVs with nonzero true coefficients (active-set RMSE) when active ASVs were randomly located (*A* = 0). Partial alignment produced consistent gains; full alignment strengthened them, particularly when taxa were rarely present. The paired PGL–PIND comparison, in which only the phylogenetic correlation differs, isolates the contribution of borrowing within each setting. Active-set size and RMS were fixed across alignment levels, although the effect distributions need not be identical. At the additional *n* = 100, *P* = 0.5, *V* = 1 setting, aligned-signal power was 0.921 for PGL and 0.895 for PIND, and the paired active-set RMSE ratio was 0.968; the gain was smaller than at *n* = 50 in the corresponding setting (Appendix F).

**Figure 1.**
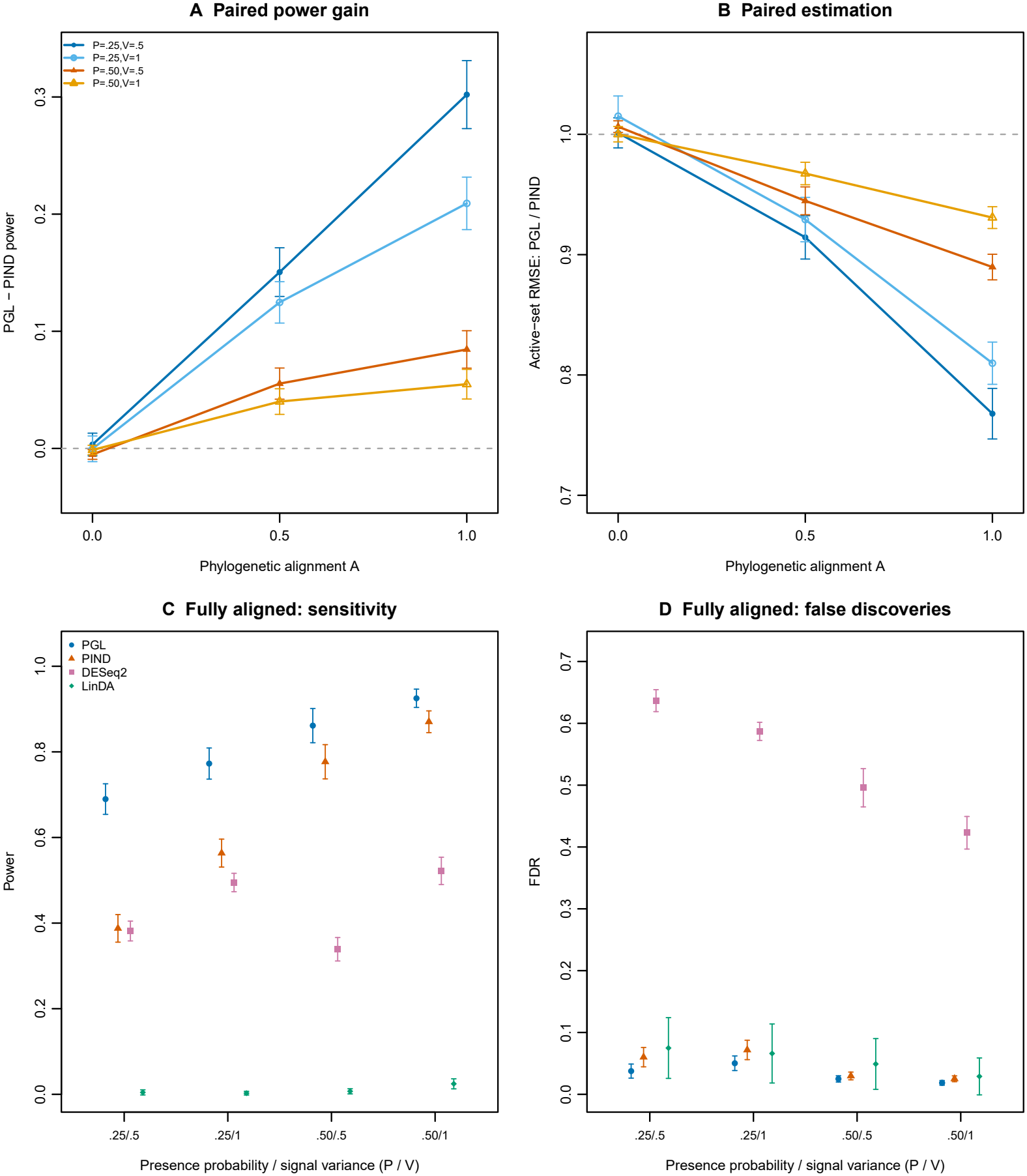
Experiment 1, *n* = 50. Panels A and B show paired PGL–PIND power differences and active-set RMSE ratios across all 12 combinations of alignment *A*, presence probability *P* , and active-effect variance *V*. Panels C and D compare power and mean FDR for PGL, PIND, DESeq2, and LinDA in all four (*P, V*) settings at full alignment (*A* = 1). Points are means over 100 replicates; bars are approximate 95% Monte Carlo intervals for those means. Data are generated from a gated multinomial logit-normal model, and Bayesian fits use the generated presence indicators. Bayesian selection uses 95% credible intervals; benchmark selection uses BH-adjusted *p <* 0.05. Ratios below one favor PGL.

The feature-wise benchmarks show why both sensitivity and false discoveries matter in this setting (Figure 1, Panels C–D). Under full alignment, PGL maintained higher power than PIND and the two benchmarks across the presence and signal-strength settings. DESeq2 detected some effects but incurred substantially more false discoveries, whereas LinDA made few discoveries under this generator. For example, in the low-presence, weaker-signal setting, PGL attained power 0.690 with mean FDR 0.038; DESeq2 attained power 0.382 with FDR 0.637. PGL’s mean FDR was also lower than PIND’s across the four full-alignment settings. These comparisons are specific to a model-based generator with tree-structured effects and known sparse support; the next experiment uses an external generator with tree-independent effects and no structural presence process.

### 3.3 Experiment 2: tree-independent external generator

Under the external generator, PGL and PIND performed similarly in selection and estimation across sample sizes and signal sparsities (Figure 2). This pattern is consistent with their agreement at *A* = 0 in Experiment 1, the direct within-generator comparison when effects were unrelated to the tree. In Experiment 2, both the count mechanism and treatment of zeros also changed, so its main role is to assess performance under a different generator and against feature-wise methods. The replicate-averaged posterior mean of *ω* decreased in the larger-sample settings, although at *n* = 50, *S* = 0.10 it remained near the 0.5 prior mean; the paired all-ASV RMSE ratio ranged from 0.999 to 1.043 across eight settings (Appendix F). Thus an unrelated tree offered little consistent benefit and could still cause modest negative transfer.

**Figure 2.**
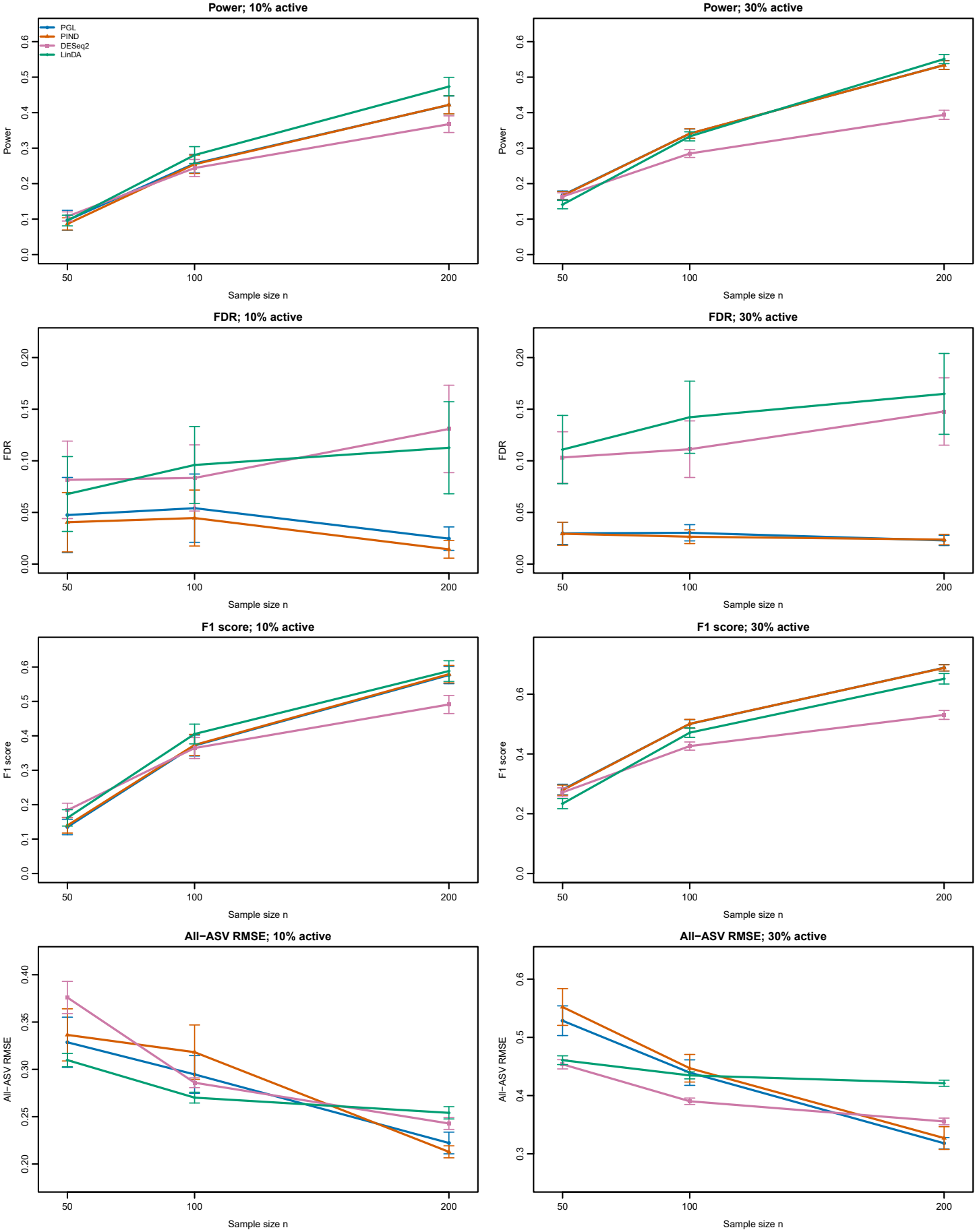
Experiment 2, tree-independent generator. Columns indicate 10% and 30% active ASVs; rows show power, mean FDR, *F*_1_, and all-ASV RMSE for PGL, PIND, DESeq2, and LinDA as sample size increases. Signal variance is *V* = 1; the *n* = 50, *V* = 0.5 settings appear in Appendix F. Points are means over 100 replicates, with approximate 95% Monte Carlo intervals. PGL and PIND curves often overlap. DESeq2 and LinDA coefficients were converted to one-unit natural-log effects while retaining their own normalization or bias correction; RMSE is measured against the common zero-sum generating effects.

The comparison with feature-wise methods changed with the performance criterion and signal sparsity. All methods gained power with sample size. LinDA could be more sensitive than PGL: at *n* = 200 it had the highest power at both active proportions. For *V* = 1 and 10% active ASVs, LinDA had the highest *F*_1_ at *n* = 100 and 200, whereas DESeq2 had the highest *F*_1_ at *n* = 50. At *n* = 200 and 10% active ASVs, PGL’s *F*_1_ was close to LinDA’s (0.577 versus 0.588). PGL and PIND had lower mean FDR than both benchmarks in every setting, under the distinct selection rules stated above. At 30% active ASVs, PGL and PIND had higher *F*_1_ at *n* = 200 despite LinDA’s slightly higher power. The plotted all-ASV RMSEs measure numerical distance to the zero-sum truth; DESeq2 and LinDA coefficients were converted to one-unit natural-log effects while retaining their own normalization or bias correction (Appendix F). These results show a sensitivity–error trade-off rather than a uniformly best method; full tables are in Appendix F.

### 3.4 Experiment 3: a misspecified input tree

The mean posterior borrowing weight responded to worsening tree information in both presence regimes (Figure 3). It was near one with the correct tree and fell steadily as tip labels were permuted, reaching about 0.2 when the supplied kernel was effectively unrelated to the generating tree. With the correct tree, adaptive PGL clearly improved power, *F*_1_, and all-ASV RMSE over PIND, especially under low presence. The adaptive model therefore retained the benefit of borrowing when the tree was useful instead of suppressing it indiscriminately. At *q* = 0, fixing *ω* = 1 gave slightly better all-ASV RMSE and *F*_1_ than learning *ω*, a small cost of adaptation when the supplied tree was correct.

**Figure 3.**
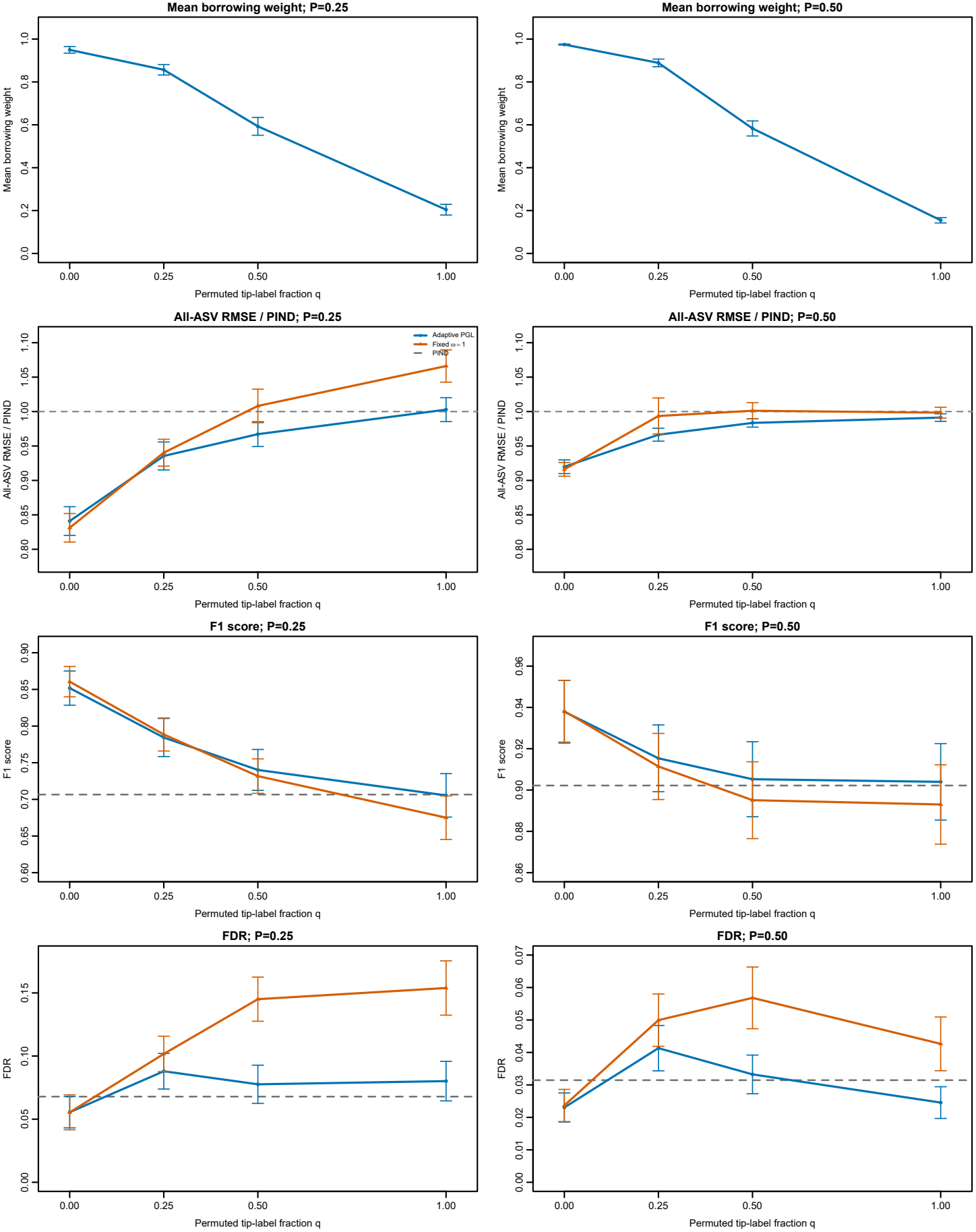
Experiment 3, corrupted input tree. Columns indicate marginal presence probabilities *P* = 0.25 and 0.50; rows show the adaptive posterior mean of *ω* averaged over replicates, all-ASV RMSE ratios to PIND, *F*_1_, and mean FDR. Adaptive and fixed-*ω* = 1 PGL are compared at each fraction *q* of permuted tip labels. Dashed lines in the lower three rows mark PIND, which does not depend on *q*; its RMSE ratio is 1. Points are means over 100 replicates with approximate 95% Monte Carlo intervals. Kernel concordance is the correlation between the unpermuted tree’s Brownian kernel and the permuted supplied kernel, falling from about 1 at *q* = 0 to 0 at *q* = 1. Experiment 3 uses independent replicates from Experiment 1.

As the tree degraded, adaptive PGL moved toward the PIND baseline, while forcing *ω* = 1 exposed a cost of borrowing based on misspecified phylogenetic relationships. At moderate corruption and low presence, adaptation kept all-ASV RMSE below PIND’s level and reduced FDR relative to forced borrowing. At complete permutation, its all-ASV RMSE was essentially at PIND’s level (ratio 1.003, Monte Carlo interval 0.985–1.020), whereas forced borrowing increased it (ratio 1.066, interval 1.043–1.090). Its *F*_1_ was likewise close to PIND’s. Its FDR was somewhat higher than PIND’s under low presence, but slightly lower under higher presence. The cost of tree error was smaller when presence was higher. Thus learning *ω* mitigated negative transfer most visibly when observations were sparse, but it did not guarantee the same selection errors as setting borrowing to zero.

## 4 Real Data Analysis

We applied PGL to salivary 16S rRNA sequencing data from the HIV study of Jayaraman et al. (2025). After removing control samples, the analysis included 26 participants living without HIV and 40 living with HIV. We retained ASVs detected in at least 10% of each group, leaving 70 ASVs, and adjusted for the clinician-assigned age group based on a 60-year cutoff. HIV status was coded as one for participants living with HIV and zero otherwise. The target coefficient describes a group-associated log-ratio change for an ASV relative to the mean across retained ASVs, conditional on each sample’s observed support; it does not estimate a group difference in ASV prevalence. The groupwise 10% filter therefore also fixes the reference for these coefficients and their interval-based selections. We used the phylogenetic tree supplied with the dataset to construct the Brownian correlation for PGL.

PGL and PIND were fitted to the same samples, ASVs, covariate design, likelihood, and priors, with PIND fixing the phylogenetic borrowing weight *ω* to zero. Four independently initialized chains per model each ran for 80,000 iterations; discarding the first 40,000 iterations and retaining every eighth subsequent draw left 20,000 combined draws. We selected an ASV when its 95% equaltailed credible interval for the group coefficient excluded zero. Appendix G gives the preprocessing, feature-level summaries, benchmark comparisons, and convergence diagnostics.

PGL and PIND yielded similar coefficient estimates but selected slightly different sets of ASVs (Figure 4). Across the 70 ASVs, PGL and PIND posterior means closely followed the identity line (Pearson correlation 0.989), and their estimated signs agreed for 94% of ASVs. All nine PIND selections were also selected by PGL; PGL selected three additional ASVs assigned to *Oribacterium, Streptococcus*, and *Scardovia*, respectively. For those three, the two methods estimated the same effect direction, but only the PGL intervals excluded zero. The posterior for *ω* suggested phylogenetic borrowing but remained broad (posterior mean 0.72, 95% credible interval 0.17–0.99; Appendix G). The structured prior shifted posterior locations and modestly altered interval widths, so that three additional ASVs met the credible-interval selection criterion under PGL while the overall effect pattern remained largely intact.

**Figure 4.**
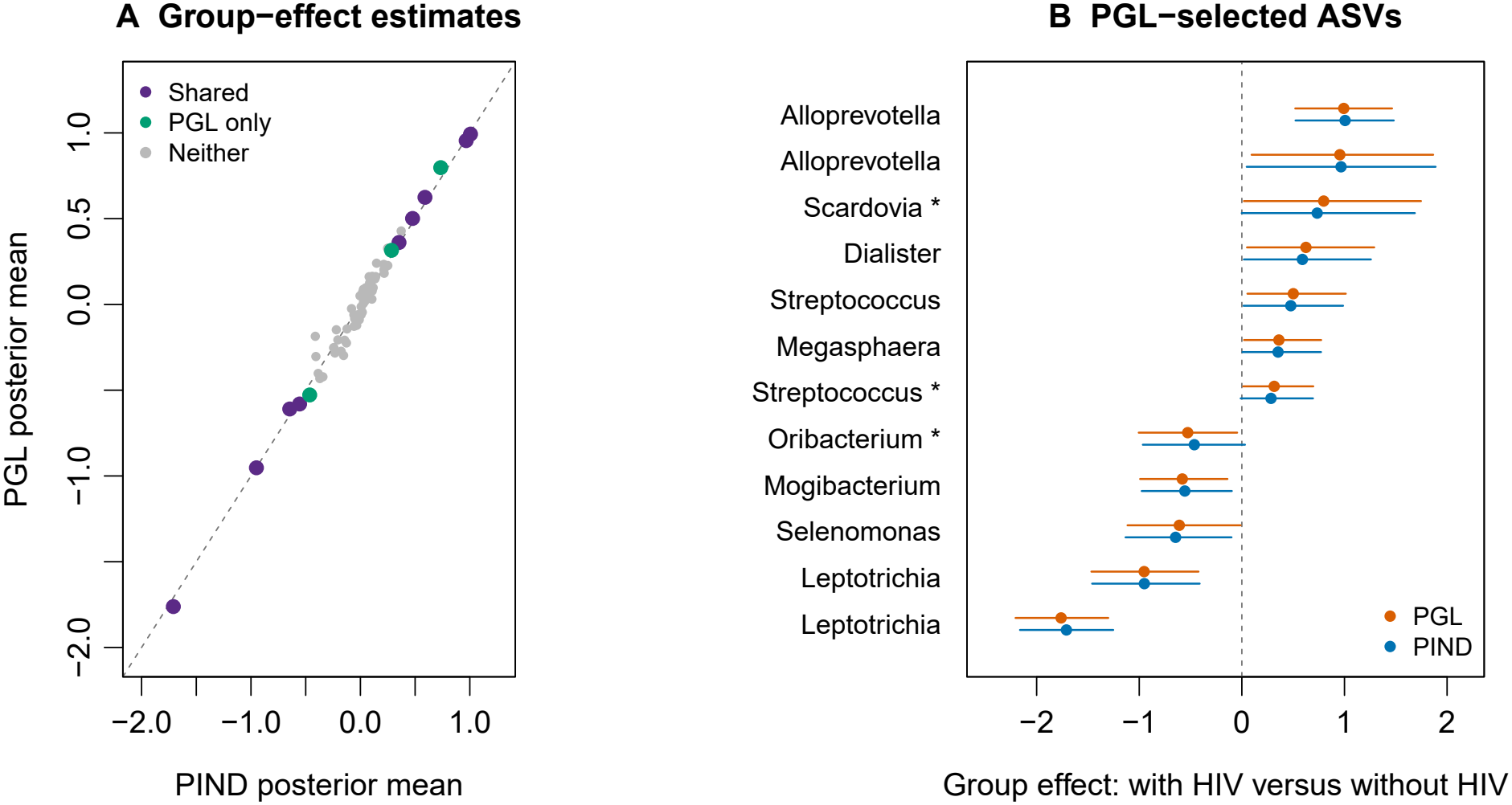
HIV oral microbiome application. Panel A compares PGL and PIND posterior means of the regression coefficients for HIV status across all 70 retained ASVs (Pearson correlation 0.989); colors distinguish shared selections, PGL-only selections, and ASVs selected by neither method. The dashed line is equality. Panel B gives posterior means and 95% equal-tailed credible intervals for the 12 ASVs selected by PGL; an asterisk marks the three selected only by PGL. Each taxonomic label denotes one ASV; repeated labels denote distinct ASVs. The dashed vertical line is zero. Positive effects indicate a larger conditional log-ratio with HIV relative to the retained-ASV reference.

The following ASV-level example illustrates the conditional abundance estimand. An ASV assigned to *Leptotrichia* was detected more frequently among participants with HIV (31/40 versus 7/26), yet its mean relative abundance among samples in which it was detected was lower (0.0042 versus 0.0148, normalized over the 70 retained ASVs). LinDA estimated a positive group coefficient for this ASV, whereas both PGL and PIND estimated negative coefficients (both posterior means −0.95; Appendix G). These opposing occurrence and positive-abundance patterns illustrate the distinction between LinDA’s zero-adjusted log-ratio regression and PGL and PIND’s log-ratio contrasts conditional on the observed support. Zero treatment and compositional reference both differ between these analyses. The negative sign also occurs under PIND without phylogenetic borrowing. At the genus level, lower *Leptotrichia* abundance in participants with HIV has been reported in an oral microbiome study spanning different levels of periodontal disease severity (Noguera-Julian et al., 2017). Complete ASV-level summaries and DESeq2 and LinDA comparisons are in Appendix G.

## 5 Discussion

PGL uses evolutionary relatedness to guide global–local shrinkage while retaining an effect for each ASV. The global scale favors sparse associations; local scales allow individual effects to remain large even when their neighbors are strongly shrunk. A Brownian correlation introduces borrowing among related standardized effects, and the exposure-specific weight *ω* lets the data moderate that borrowing. The zero-sum constraint gives each coefficient a log-ratio interpretation relative to the mean across retained ASVs. Thus phylogeny informs feature-specific inference without requiring taxonomic aggregation. PIND, obtained by fixing *ω* = 0 within the same likelihood and prior construction, isolates the contribution of structured borrowing.

The simulations show where that contribution is most useful: related ASVs can support one another when effects align with the tree and individual observations are sparse. The tree-independent external generator yielded similar PGL and PIND results, while the feature-wise benchmarks exhibited different sensitivity–error trade-offs under distinct selection rules. When the supplied tree was corrupted, learning *ω* reduced the harm from forced borrowing without making selection identical to PIND. These findings support adaptive use of the supplied structure, with the strength and precision of borrowing still dependent on the data and the tree.

In the HIV oral microbiome analysis, PGL and PIND agreed closely on effect directions and magnitudes, while borrowing shifted posterior locations and interval endpoints for a few ASVs. The *Leptotrichia* ASV comparison further shows why the observed-support target matters when interpreting results alongside methods that adjust zeros before log-ratio transformation.

The interpretation of those effects depends on the observed-support gate. It conditions the multinomial abundance model on detected ASVs; it does not model their occurrence probabilities or distinguish biological absence from sequencing and technical zeros. Accordingly, a regression co-efficient for HIV status is a conditional log-ratio association relative to the retained-ASV reference, whereas marginal or pseudocount-based analyses can combine occurrence and positive-abundance differences. A joint prevalence–abundance model would make both components explicit. Phylogenetic borrowing in a prevalence layer is biologically plausible, but co-occurrence of related taxa does not by itself establish shared exposure effects (Faust et al., 2012; Martiny et al., 2015). An extension could therefore use separate kernels for prevalence and abundance, with an estimable cross-process association that includes independence. Repeated samples, technical controls, or absolute-abundance measurements would help distinguish absence from non-detection and identify that additional structure.

The analysis also treats the tree-derived correlation matrix as fixed and does not account for uncertainty in the underlying tree. Marker-gene pipelines commonly infer an unrooted tree and then place a working root; for example, the widely used QIIME 2 MAFFT–FastTree pipeline supplies a midpoint-rooted tree (QIIME 2 Development Team, 2026). In the HIV oral microbiome analysis, we retained the existing root of the phylogenetic tree relating the microbial ASVs. The root location can affect the Brownian shared-path correlation, but the tree-corruption experiment varied tip assignments, not rooting. Studying alternative rooting conventions, along with topology, branch-length, marker-gene, and kernel uncertainty, is a direction for future work rather than part of the current model comparison. Credible-interval exclusion is a transparent selection rule but does not provide a formal false-discovery-rate guarantee; calibrated posterior decision rules and replication in independent cohorts are needed for stronger discovery claims.

Finally, the sampler was practical at the dimensions studied. In the HIV oral microbiome analysis, four independently initialized MCMC chains for each model yielded consistent posterior summaries of the regression coefficients for HIV status (Appendix G). Dense Gaussian operations for the effect vector and sample-by-feature latent abundance updates will limit scaling as the number of ASVs grows. Faster tree-based factorizations and more efficient block updates are natural computational directions. PGL therefore provides a feature-specific framework for adaptive phylogenetic borrowing, with its gains, interpretation, and limitations tied to the information carried by the tree and the observed support.

## Acknowledgements

This work was supported by the School of Medicine, Case Western Reserve University [BGT630267 to L.Z.]. OpenAI’s Codex was used to assist with manuscript editing and code revision; the authors take responsibility for the final content.

## Data Availability

The 16S rRNA sequencing data used in the HIV oral microbiome analysis were reported by Jayaraman et al. (2025) and are publicly available in Qiita under study ID 16115 (https://qiita.ucsd.edu/study/description/16115).

## Appendices

### A Brownian Phylogenetic Correlation and Adaptive Kernel

#### A.2 Tip correlation from a rooted tree

Let *T* be a rooted tree whose tips are the *K* retained ASVs, in the same order as the columns of the count table. The implementation requires branch lengths. If an edge length is nonfinite or less than 10^−6^ times the smallest positive edge length, it is replaced by that floor. For tips *k* and *h*, define the Brownian covariance

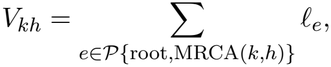

where *P{*root, MRCA(*k, h*)*}* is the shared root-to-ancestor path and *l*_*e*_ is the length of edge *e*. In particular, *V*_*kk*_ is the root-to-tip length. This is the covariance computed by `ape::vcv.phylo` with its Brownian model. After matching the tip order, we form

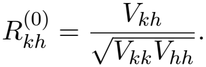

The off-diagonal entries measure shared evolutionary history relative to the total history of each tip; they are generally nonnegative. Their sign does not prescribe the sign of the final compositional coefficient covariances.

For numerical stability, the implementation symmetrizes ***R***^(0)^, floors its eigenvalues at 10^−8^, reconstructs the matrix, and rescales it to unit diagonal. If its smallest eigenvalue is still below 10^−8^, it adds the minimum identity mixture needed to reach that floor. We denote the resulting positive-definite correlation matrix by ***R***_phylo_. These small adjustments are numerical, rather than a different phylogenetic kernel.

#### A.2 Borrowing strength

For exposure group *g*,

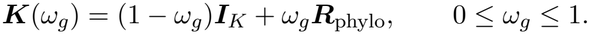

Because both component matrices have unit diagonal, *K*_*kk*_(*ω*_*g*_) = 1. The off-diagonal entries are *ω*_*g*_*R*_phylo,*kh*_, so *ω*_*g*_ controls dependence among the standardized latent effects without multiplying their marginal variance. For 0 *≤ ω*_*g*_ *≤* 1, ***K***(*ω*_*g*_) is positive definite. The prior is *ω*_*g*_ *∼* Beta(1, 1) independently across exposure groups. All non-reference indicators of one categorical exposure may share an *ω*_*g*_; continuous exposures ordinarily receive their own. PIND fixes every *ω*_*g*_ = 0 and otherwise uses the same likelihood, projection, shrinkage priors, and inference algorithm.

### B Dependence after Global–Local Scaling and Projection

#### B.1 Coefficient covariance

For exposure column *j*, let *g*(*j*) be its exposure group and write

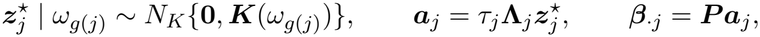

where 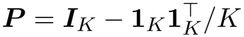 and **Λ**_*j*_ = diag(*λ*_1*j*_, … , *λ*_*Kj*_). Conditional on the scales and *ω*_*g*(*j*)_,

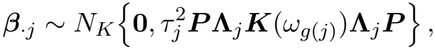

as a Gaussian distribution on the (*K −* 1)-dimensional contrast space. The matrix is singular in the direction **1**_*K*_, exactly as required by 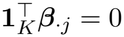. Elementwise,

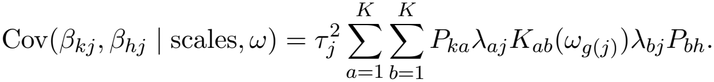

It is therefore incorrect to use 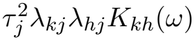 as the covariance of the final *β* coordinates; that expression applies to the unprojected ***a***_*j*_. Even when *ω* = 0,

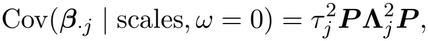

which generally has nonzero off-diagonal entries. Thus PIND removes phylogenetic dependence before projection, but retains dependence induced by the zero-sum contrast. Changing one *λ*_*kj*_ so that the unprojected *a*_*kj*_ changes by Δ, while holding the other latent coordinates fixed, changes *β*_*kj*_ by (1 − 1*/K*)Δ and each *β*_*hj*_, *h ≠ k*, by −Δ*/K*. The local scale permits a feature-specific departure; it does not create an exactly sparse ***β*** or leave the other coefficients unchanged.

#### B.2 Latent pseudo-abundance covariance

For sample *i*, condition on ***η***_*i*_, the scales, the borrowing parameters, **Γ**, and *σ*_*i*_. Let *c*_*ik*_ = *η*_*ik*_ exp(*u*_*ik*_) and

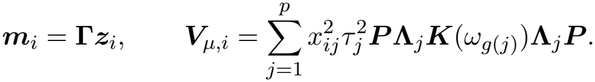

The latent effect vectors are conditionally independent across exposure columns, even when some share *ω*_*g*_. Integrating over them yields 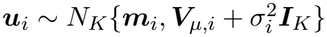. Consequently, for *k≠ h*,

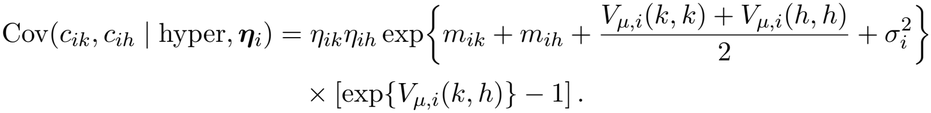

This follows from the multivariate-normal moment-generating function. The covariance is zero when either gate is zero. Otherwise its sign is the sign of the *projected* covariance *V*_*µ,i*_(*k, h*), which can be negative despite nonnegative Brownian tip correlations. This is a covariance of latent pseudo-abundances under the prior, not a covariance of observed multinomial counts.

### C Support Conditioning and Contrast Interpretation

#### C.1 What the gate conditions on

The gate *η*_*ik*_ = *I*(*y*_*ik*_ *>* 0) is computed from the observed count table and then held fixed. The multinomial factor in the fitted model is

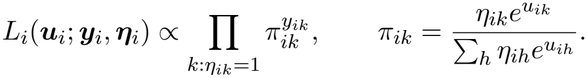

This is a working likelihood on the observed support. It does not assign a probability to ***η***_*i*_, and it does not include the normalizing factor that an exact multinomial distribution truncated to positive counts in every active category would require. Thus it should not be interpreted as a joint generative model for detection and abundance. In particular, a zero can reflect biological absence, sequencing depth, or technical variation, and the model does not determine which occurred. The Gaussian layer includes a latent *u*_*ik*_ even when *η*_*ik*_ = 0; that coordinate has no direct multinomial contribution in sample *i*.

#### C.2 Likelihood invariance and the chosen reference

For any scalar *a*_*i*_, replacing all active *u*_*ik*_ by *u*_*ik*_ + *a*_*i*_ leaves ***π***_*i*_ unchanged. If *η*_*ik*_ = *η*_*ih*_ = 1, then

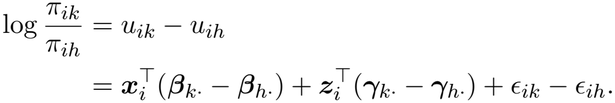

Thus, at fixed observed support and residuals, the slope for a one-unit change in *x*_*ij*_ is the pairwise log-ratio contrast *β*_*kj*_ *− β*_*hj*_. An unrestricted common shift of all the *β*_*kj*_ for an exposure would not change these contrasts and would be prior-anchored. We instead impose the exact constraint ∑_*k*_ *β*_*kj*_ = 0, giving

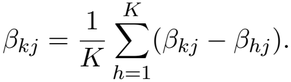

The coefficient is therefore a slope relative to the unweighted mean effect across the *K* retained ASVs. Changing the set of retained ASVs changes this reference. It is not the slope of absolute abundance, and the nonlinear multinomial normalization means it is not generally a marginal change in expected relative abundance either.

An equal-tailed credible interval for *β*_*kj*_ that excludes zero supports a nonzero contrast relative to this reference under the fitted model. It does not assert that every pairwise contrast is nonzero, that the ASV’s prevalence changes, or that its biological absolute abundance changes. The exact zero-sum constraint removes the common exposure-effect direction. Sparse co-observation, collinear covariates, and the fixed support may still leave individual contrasts weakly informed; the global– local prior regularizes them. Nuisance coefficient vectors are not projected and retain proper Gaussian priors. For categorical exposures, all non-reference indicators enter together; a contrast between two non-reference levels is a difference of their coefficient vectors and also sums to zero.

### D Prior Specification and Calibration

#### D.1 Shrinkage and borrowing priors

The independent global and local scales have density

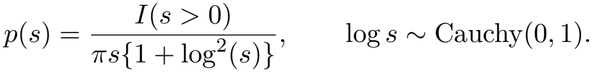

Its pole near zero encourages shrinkage, while its upper tail permits large latent departures (Zhu and Tang, 2025). After projection, these departures are interpreted as contrasts, and neither *s* nor *β*_*kj*_ has a point mass at zero. The borrowing parameters use independent Beta(1, 1) priors. The log-Cauchy and Brownian Gaussian components address different roles: the scales control magnitude and heterogeneity, while *ω*_*g*_ controls pre-projection phylogenetic correlation.

#### D.2 Nuisance coefficients and residual scales

Independently,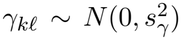, including an intercept coefficient when an intercept is in ***z***_*i*_. To place the central 95% prior range for exp(*γ*_*kl*_) within [1*/r, r*] for a one-unit covariate change, choose *s*_*γ*_ = log(*r*)*/*1.96. Taking *r* = 4 gives *s*_*γ*_ *≃* 0.71. This is a prior calibration for a latent log-weight coefficient, not a guarantee about a marginal abundance ratio.

The sample-specific *σ*_*i*_ has a half-normal prior with scale *s*_*σ*_. Since the residual difference of two latent log weights has standard deviation 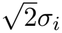, a reference 95% *r*-fold contrast corresponds to

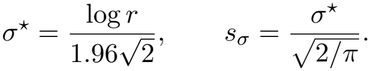

With *r* = 4, *s*_*σ*_ *≃* 0.63. These are the default scales used in the implementation.

### E Posterior Computation

#### E.2 Notation and Gaussian blocks

Write ***u***_*i*_ = (*u*_*i*1_, … , *u*_*iK*_)^*⊤*^ and ***µ***_*i*_ = ∑_*j*_ *x*_*ij*_***β***_*·j*_ + **Γ*z***_*i*_. For exposure column *j*, let

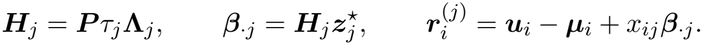

Conditional on the other blocks, 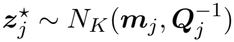, with

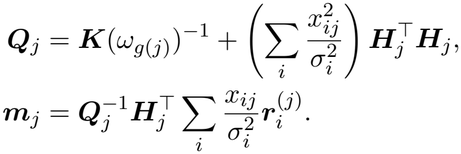

The projection is inside ***H***_*j*_; omitting it would give the full conditional for the unconstrained model. A Cholesky factorization of ***Q***_*j*_ is used for the block draw. For nuisance column *l*, set 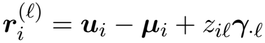. Then ***γ***_*·l*_ *∼ N*_*K*_(*v*_*l*_***b***_*l*_, *v*_*l*_***I***_*K*_), where

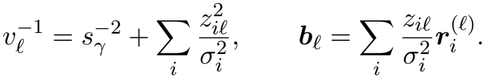

#### E.2 Latent log weights

Let 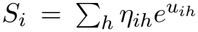. For coordinate (*i, k*), propose 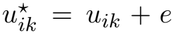 with 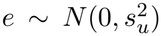, and set 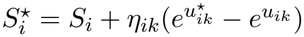. The Metropolis log ratio is

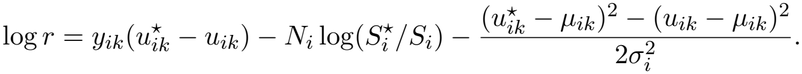

For *η*_*ik*_ = 0, *y*_*ik*_ = 0 and the multinomial terms vanish; the Gaussian prior term still updates that latent coordinate.

#### E.3 Global and local scale updates

For *τ*_*j*_, define 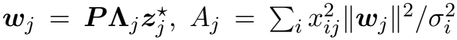, and 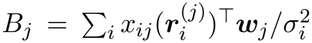. Propose a Gaussian random walk in *a*_*j*_ = log *τ*_*j*_. The log Metropolis ratio is

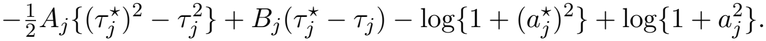

The final two terms combine the positive log-Cauchy density with the Jacobian of the log-scale proposal.

For 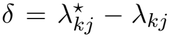 and 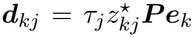, where ***e***_*k*_ is the *k*th coordinate vector. The proposed coefficient change is *δ****d***_*kj*_ across *all K* coordinates. With ***e***_*i*_ = ***u***_*i*_ − ***µ***_*i*_ at the current state, the log likelihood ratio is

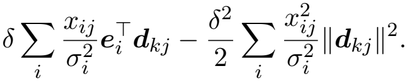

A Gaussian random walk in *b*_*kj*_ = log *λ*_*kj*_ adds the transformed log-Cauchy prior ratio 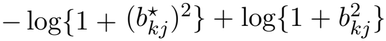. After these updates the implementation recomputes 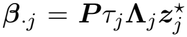 preserving the exact zero-sum constraint.

#### E.4 Borrowing and residual-scale updates

The eigendecomposition ***R***_phylo_ = ***U*** diag(*r*_1_, … , *r*_*K*_)***U*** ^*⊤*^ is cached. The eigenvalues of ***K***(*ω*) are 1 *− ω* + *ωr*_*h*_, allowing its determinant and inverse to be evaluated for each proposal. For group *g*, propose a Gaussian random walk in *q*_*g*_ = logit(*ω*_*g*_). The Metropolis log ratio is

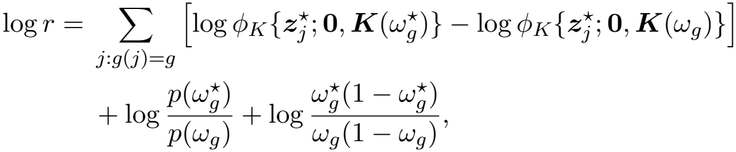

where *p* is the Beta(1, 1) density. The last term is the logit-transformation Jacobian. PIND fixes *ω*_*g*_ = 0 and omits this update.

For sample *i*, let *Q*_*i*_ = *∥****u***_*i*_ − ***µ***_*i*_*∥*^2^ over all *K* latent coordinates. Propose a Gaussian random walk in *c*_*i*_ = log *σ*_*i*_. Under the half-normal prior, the log ratio is

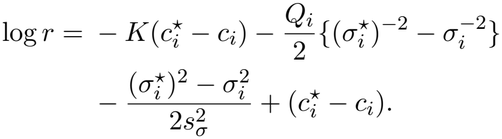

The gate affects the multinomial likelihood, not the Gaussian residual sum *Q*_*i*_.

Proposal scales are adapted during warm-up toward an acceptance rate of 0.40 and then held fixed. Posterior inference uses the stored, projected ***β*** draws. The primary selection rule excludes zero from a 95% equal-tailed credible interval for the relevant coefficient or prespecified group contrast. Markov chain Monte Carlo (MCMC) assessment focuses on these target contrasts rather than on individual shrinkage scales.

### F Complete Simulation Designs and Results

#### F.1 Experiment 1: tree-aligned signals

For each replicate we generated a 200-tip random tree. The data generator formed a tip correlation ***C***_DGP_ from a standardized pseudoinverse of its normalized graph Laplacian, with a stabilizing ridge if required. This correlation governed a Gaussian-probit presence vector in each sample, thresholded at the requested marginal presence probability *P*. The *fitted* PGL prior instead used the Brownian tip correlation ***R***_phylo_ from the same tree (Appendix A); these are distinct tree-based matrices. If a generated sample had no present ASVs, one tip was set present to define the multinomial probability.

Exactly 60 of 200 exposure coefficients were active. Alignment *A ∈ {*0, 0.5, 1*}* assigned 0, 30, or 60 active ASVs to nonoverlapping descendant-tip groups of size 5–20; remaining active ASVs were selected without reference to the tree. Each clade had a nonzero random effect center and correlated within-clade deviations of scale 0.10 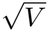, using the corresponding submatrix of ***C***_DGP_; unstructured effects were generated separately. We subtracted the mean of the *active* effects and rescaled their RMS to 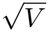. Consequently the final target has exactly 60 nonzero entries, 140 exact zeros, zero total sum, and the same RMS of the nonzero true coefficients in every alignment setting. This correction avoids treating inactive ASVs as nonzero merely because an initially sparse vector was centered across all 200 ASVs.

One target exposure and five nuisance covariates were independent standard normal variates. Nuisance coefficients were *N* (0, 0.3^2^), and sample-specific residual standard deviations were uniform on (0.4, 0.6). We generated 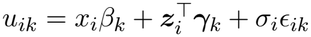, with independent standard normal *ϵ*_*ik*_, and drew 50,000 multinomial reads per sample with probabilities proportional to 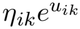. The Bayesian fits used the generated presence indicators *η*_*ik*_. A present ASV that received zero reads remained in the multinomial denominator. The 12 main settings crossed *n* = 50, three values of *A*, two values of *P* , and *V ∈ {*0.5, 1*}*. Two targeted sample-size settings used *n* = 100, *P* = 0.5, *V* = 1, and *A ∈ {*0, 1*}*. All 14 settings have 100 completed replicates.

**Table S1:**
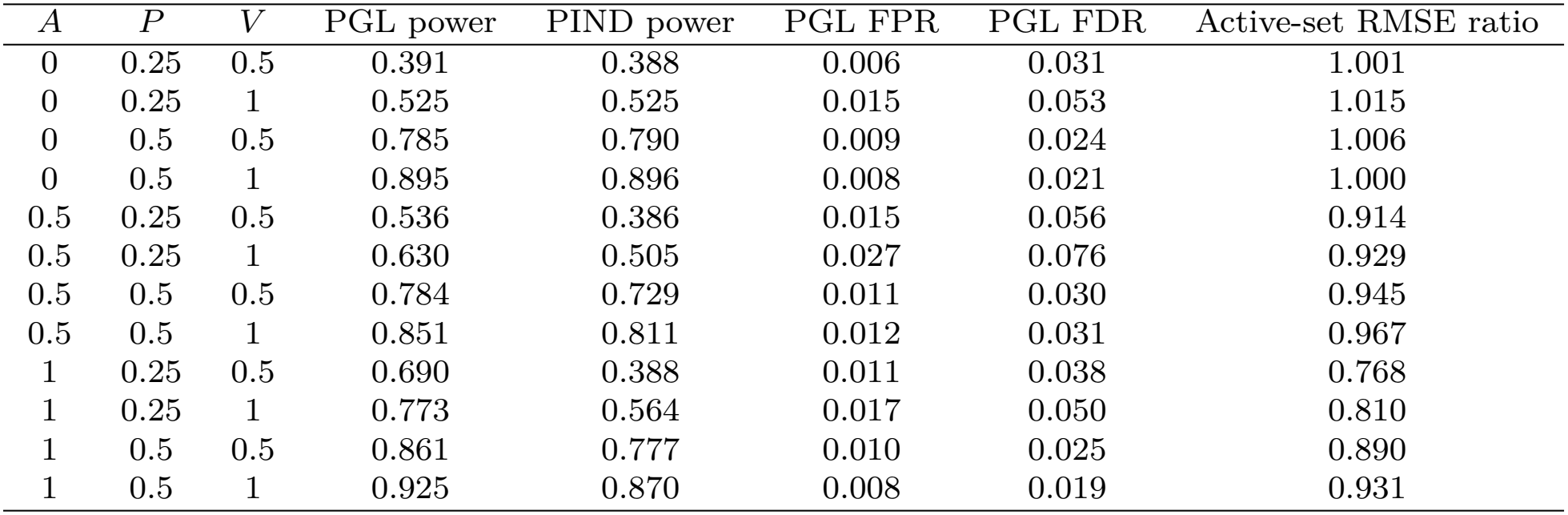
Experiment 1, *n* = 50: Monte Carlo means over 100 replicates per setting. PGL and PIND power use 95% credible intervals. The root mean squared error (RMSE) column is the mean paired active-set PGL/PIND ratio.

| *A* | *P* | *V* | PGL power | PIND power | PGL FPR | PGL FDR | Active-set RMSE ratio |
|---|---|---|---|---|---|---|---|
| 0 | 0.25 | 0.5 | 0.391 | 0.388 | 0.006 | 0.031 | 1.001 |
| 0 | 0.25 | 1 | 0.525 | 0.525 | 0.015 | 0.053 | 1.015 |
| 0 | 0.5 | 0.5 | 0.785 | 0.790 | 0.009 | 0.024 | 1.006 |
| 0 | 0.5 | 1 | 0.895 | 0.896 | 0.008 | 0.021 | 1.000 |
| 0.5 | 0.25 | 0.5 | 0.536 | 0.386 | 0.015 | 0.056 | 0.914 |
| 0.5 | 0.25 | 1 | 0.630 | 0.505 | 0.027 | 0.076 | 0.929 |
| 0.5 | 0.5 | 0.5 | 0.784 | 0.729 | 0.011 | 0.030 | 0.945 |
| 0.5 | 0.5 | 1 | 0.851 | 0.811 | 0.012 | 0.031 | 0.967 |
| 1 | 0.25 | 0.5 | 0.690 | 0.388 | 0.011 | 0.038 | 0.768 |
| 1 | 0.25 | 1 | 0.773 | 0.564 | 0.017 | 0.050 | 0.810 |
| 1 | 0.5 | 0.5 | 0.861 | 0.777 | 0.010 | 0.025 | 0.890 |
| 1 | 0.5 | 1 | 0.925 | 0.870 | 0.008 | 0.019 | 0.931 |

**Table S2:** Experiment 1, *n* = 50: PIND false-positive rate (FPR), mean false discovery rate (FDR), and *F*_1_ under the 95% credible-interval selection rule. Entries are Monte Carlo means over 100 replicates per setting and complete the PIND comparison in Table S1.

| *A* | *P* | *V* | FPR | FDR | *F*<sub>1</sub> |
|---|---|---|---|---|---|
| 0 | 0.25 | 0.5 | 0.005 | 0.023 | 0.548 |
| 0 | 0.25 | 1 | 0.009 | 0.035 | 0.674 |
| 0 | 0.5 | 0.5 | 0.008 | 0.023 | 0.871 |
| 0 | 0.5 | 1 | 0.008 | 0.020 | 0.935 |
| 0.5 | 0.25 | 0.5 | 0.008 | 0.041 | 0.536 |
| 0.5 | 0.25 | 1 | 0.012 | 0.051 | 0.649 |
| 0.5 | 0.5 | 0.5 | 0.007 | 0.023 | 0.824 |
| 0.5 | 0.5 | 1 | 0.009 | 0.024 | 0.877 |
| 1 | 0.25 | 0.5 | 0.012 | 0.060 | 0.526 |
| 1 | 0.25 | 1 | 0.021 | 0.072 | 0.684 |
| 1 | 0.5 | 0.5 | 0.011 | 0.030 | 0.844 |
| 1 | 0.5 | 1 | 0.010 | 0.025 | 0.914 |

At *n* = 100, the active-set RMSE ratios were 1.000 at *A* = 0 and 0.968 at *A* = 1. At *A* = 1, mean power was 0.921 for PGL and 0.895 for PIND; at *A* = 0 the corresponding means were 0.980 and 0.980. The comparison used all 100 replicates in each setting. The completed run also included DESeq2 and LinDA at every Experiment 1 setting. DESeq2 used the target exposure and five nuisance covariates, local dispersion, and a unit pseudo-count for size-factor estimation in the structurally sparse generator; LinDA used its explicit pseudo-count zero handler. Their selection rules were BH-adjusted *p <* 0.05, while Bayesian methods used the credible-interval rule described in the main text. In this generator the feature-wise procedures often had appreciably higher FDR or very low power; this motivated the tree-independent generator in Experiment 2 rather than a claim of general dominance from a model-based experiment.

**Table S3:** Experiment 1, *n* = 50: feature-wise benchmark selection under BH-adjusted *p <* 0.05. Entries are Monte Carlo means over 100 replicates per setting.

| *A* | *P* | *V* | DESeq2 power | DESeq2 FDR | DESeq2 *F*<sub>1</sub> | LinDA power | LinDA FDR | LinDA *F*<sub>1</sub> |
|---|---|---|---|---|---|---|---|---|
| 0 | 0.25 | 0.5 | 0.416 | 0.607 | 0.398 | 0.003 | 0.056 | 0.004 |
| 0 | 0.25 | 1 | 0.509 | 0.573 | 0.460 | 0.003 | 0.059 | 0.005 |
| 0 | 0.5 | 0.5 | 0.344 | 0.477 | 0.403 | 0.004 | 0.063 | 0.007 |
| 0 | 0.5 | 1 | 0.490 | 0.412 | 0.525 | 0.013 | 0.047 | 0.024 |
| 0.5 | 0.25 | 0.5 | 0.401 | 0.619 | 0.384 | 0.005 | 0.047 | 0.007 |
| 0.5 | 0.25 | 1 | 0.496 | 0.582 | 0.449 | 0.004 | 0.057 | 0.006 |
| 0.5 | 0.5 | 0.5 | 0.352 | 0.473 | 0.408 | 0.006 | 0.059 | 0.011 |
| 0.5 | 0.5 | 1 | 0.475 | 0.425 | 0.509 | 0.013 | 0.052 | 0.024 |
| 1 | 0.25 | 0.5 | 0.382 | 0.637 | 0.367 | 0.004 | 0.075 | 0.007 |
| 1 | 0.25 | 1 | 0.495 | 0.587 | 0.445 | 0.003 | 0.066 | 0.004 |
| 1 | 0.5 | 0.5 | 0.339 | 0.496 | 0.392 | 0.007 | 0.049 | 0.012 |
| 1 | 0.5 | 1 | 0.522 | 0.423 | 0.532 | 0.025 | 0.029 | 0.042 |

#### F.2 Experiment 2: tree-independent external generator

This generator followed a LinDA-inspired log-normal absolute-abundance and multinomial sampling scheme. A binary exposure was drawn with probability logit^−1^(0.5*z*_*i*1_+0.5*z*_*i*2_), where *z*_*i*1_ is a random *{−*1, 1*}* covariate and *z*_*i*2_ is standard normal. Both were adjusted in all fitted methods. Active ASVs were chosen independently of a random tree, with balanced positive and negative effects; centering and rescaling within the active set gave exact zero sum, exact null effects, and RMS of the nonzero true coefficients 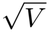. A randomly permuted Zipf rank-abundance curve formed the ASV intercepts. Taxon-specific Gaussian residual standard deviations were uniform on (0.5, 1), and two nuisance-effect vectors were generated from *N* (0, 0.3^2^). The exponentiated abundances were normalized before sampling; library sizes followed a negative-binomial law with mean 7,645 and size 5.3, bounded below by 1,000. All taxa were structurally present, so count zeros were sampling zeros and the Bayesian fits set *η*_*ik*_ = 1, including for observed zero counts. The random tree was used only for fitting PGL.

At *n* = 50, active proportions 0.10 and 0.30 were crossed with *V ∈ {*0.5, 1*}*. At *n* = 100 and 200, both active proportions were run with *V* = 1. All eight settings have 100 completed replicates. DESeq2 used poscounts size factors for this sampling-zero generator, and LinDA used its pseudo-count zero handler. DESeq2 coefficients were multiplied by log 2; LinDA coefficients were divided by the exposure standard deviation and multiplied by log 2. These conversions put them on a one-unit natural-log scale. The benchmark coefficient vectors retain their own normalization or bias correction and were not recentered. Cross-method RMSEs measure numerical distance to the common zero-sum generating effects and include any estimated common shift as well as errors in the contrasts.

**Table S4:** Experiment 2: mean power and mean FDR over 100 replicates per setting. Bayesian methods use 95% credible intervals; feature-wise benchmarks use BH-adjusted *p <* 0.05. Method columns follow the same order in both blocks.

|  |  |  | Power |  |  |  | FDR |  |  |  |
|---|---|---|---|---|---|---|---|---|---|---|
| *n* | *S* | *V* | PGL | PIND | DESeq2 | LinDA | PGL | PIND | DESeq2 | LinDA |
| 50 | 0.1 | 0.5 | 0.012 | 0.013 | 0.038 | 0.026 | 0.008 | 0.003 | 0.070 | 0.085 |
| 50 | 0.1 | 1 | 0.097 | 0.086 | 0.107 | 0.096 | 0.047 | 0.041 | 0.082 | 0.068 |
| 50 | 0.3 | 0.5 | 0.062 | 0.049 | 0.066 | 0.052 | 0.041 | 0.027 | 0.078 | 0.083 |
| 50 | 0.3 | 1 | 0.168 | 0.166 | 0.164 | 0.141 | 0.030 | 0.029 | 0.103 | 0.111 |
| 100 | 0.1 | 1 | 0.257 | 0.255 | 0.244 | 0.281 | 0.054 | 0.045 | 0.083 | 0.096 |
| 100 | 0.3 | 1 | 0.341 | 0.341 | 0.285 | 0.333 | 0.030 | 0.026 | 0.111 | 0.142 |
| 200 | 0.1 | 1 | 0.421 | 0.422 | 0.367 | 0.473 | 0.025 | 0.014 | 0.131 | 0.113 |
| 200 | 0.3 | 1 | 0.534 | 0.534 | 0.394 | 0.551 | 0.023 | 0.024 | 0.148 | 0.165 |

**Table S5:** Experiment 2: mean *F*_1_ scores across 100 replicates per setting, using each method’s primary selection rule.

| *n* | *S* | *V* | PGL | PIND | DESeq2 | LinDA |
|---|---|---|---|---|---|---|
| 50 | 0.1 | 0.5 | 0.021 | 0.025 | 0.070 | 0.047 |
| 50 | 0.1 | 1 | 0.135 | 0.140 | 0.184 | 0.162 |
| 50 | 0.3 | 0.5 | 0.086 | 0.079 | 0.119 | 0.094 |
| 50 | 0.3 | 1 | 0.281 | 0.279 | 0.272 | 0.234 |
| 100 | 0.1 | 1 | 0.372 | 0.374 | 0.365 | 0.405 |
| 100 | 0.3 | 1 | 0.500 | 0.501 | 0.426 | 0.471 |
| 200 | 0.1 | 1 | 0.577 | 0.579 | 0.491 | 0.588 |
| 200 | 0.3 | 1 | 0.688 | 0.688 | 0.531 | 0.652 |

**Table S6:** Experiment 2: mean all-ASV RMSE for each method, mean paired PGL/PIND RMSE ratio, and mean posterior borrowing weight. Each setting has 100 replicates. The paired ratio averages replicate-level ratios and need not equal the ratio of the displayed mean RMSEs. Benchmark coefficients were converted to natural-log units but retain their own reference, so cross-method absolute RMSEs are descriptive.

| *n* | *S* | *V* | PGL RMSE | PIND RMSE | DESeq2 RMSE | LinDA RMSE | Paired ratio | Mean *ω* |
|---|---|---|---|---|---|---|---|---|
| 50 | 0.1 | 0.5 | 0.231 | 0.230 | 0.344 | 0.259 | 1.003 | 0.503 |
| 50 | 0.1 | 1 | 0.329 | 0.336 | 0.376 | 0.310 | 1.023 | 0.484 |
| 50 | 0.3 | 0.5 | 0.429 | 0.439 | 0.388 | 0.353 | 1.021 | 0.444 |
| 50 | 0.3 | 1 | 0.529 | 0.552 | 0.454 | 0.461 | 1.000 | 0.293 |
| 100 | 0.1 | 1 | 0.295 | 0.318 | 0.286 | 0.270 | 0.999 | 0.419 |
| 100 | 0.3 | 1 | 0.439 | 0.447 | 0.390 | 0.435 | 1.027 | 0.209 |
| 200 | 0.1 | 1 | 0.222 | 0.213 | 0.243 | 0.254 | 1.043 | 0.373 |
| 200 | 0.3 | 1 | 0.318 | 0.327 | 0.356 | 0.421 | 1.000 | 0.167 |

Table S6 gives the estimation comparison. The observed zero fraction averaged about 0.66 in each setting. The signal and tree were generated independently. The paired PGL/PIND ratios in Table S6 quantify the small estimation costs possible with an uninformative tree; they do not imply uniform superiority over either feature-wise benchmark.

#### F.3 Experiment 3: input-tree misspecification

Data were generated as in Experiment 1 with *n* = 50, *K* = 200, *A* = 1, *V* = 1, and *P ∈ {*0.25, 0.50*}*. The Bayesian fits used the generated presence indicators as in Experiment 1. The true tree remained fixed within a replicate. For each corruption fraction *q ∈ {*0, 0.25, 0.5, 1*}*, a fraction *q* of tip labels was selected and permuted in the Brownian correlation supplied to the fit. The data and true coefficients did not change with *q*. Adaptive PGL and forced-*ω* = 1 PGL were fitted at all four levels; PIND was fitted once per replicate and presence regime. Each of the two regimes contains 100 complete paired replicates. The realized correlation between upper triangles of the true and fitted kernels averaged about 1.00, 0.61, 0.30, and 0.00 for *q* = 0, 0.25, 0.5, 1, respectively.

**Table S7:** Experiment 3: Monte Carlo means over 100 replicates per presence regime. All-ASV RMSE ratios use the same paired PIND baseline; FDR and *F*_1_ use 95% credible intervals. “Forced” fixes *ω* = 1.

| *P* | *q* | Adaptive<br>ω | Adaptive<br>RMSE ratio | Forced<br>RMSE ratio | Adaptive<br>FDR | Forced<br>FDR | Adaptive<br>*F*<sub>1</sub> | Forced<br>*F*<sub>1</sub> |
|---|---|---|---|---|---|---|---|---|
| 0.25 | 0 | 0.950 | 0.841 | 0.831 | 0.056 | 0.055 | 0.852 | 0.861 |
| 0.25 | 0.25 | 0.857 | 0.936 | 0.940 | 0.088 | 0.102 | 0.784 | 0.789 |
| 0.25 | 0.5 | 0.593 | 0.967 | 1.008 | 0.078 | 0.145 | 0.740 | 0.732 |
| 0.25 | 1 | 0.204 | 1.003 | 1.066 | 0.080 | 0.154 | 0.705 | 0.675 |
| 0.5 | 0 | 0.974 | 0.920 | 0.916 | 0.023 | 0.024 | 0.938 | 0.938 |
| 0.5 | 0.25 | 0.889 | 0.966 | 0.994 | 0.041 | 0.050 | 0.915 | 0.911 |
| 0.5 | 0.5 | 0.583 | 0.984 | 1.001 | 0.033 | 0.057 | 0.905 | 0.895 |
| 0.5 | 1 | 0.155 | 0.991 | 0.998 | 0.025 | 0.043 | 0.904 | 0.893 |

The posterior mean *ω* decreased strictly across all four corruption levels in 86% of the *P* = 0.25 replicates and 91% of the *P* = 0.50 replicates. The PIND baseline had mean power, FDR, and *F*_1_ of 0.592, 0.068, and 0.707 at *P* = 0.25, and 0.858, 0.031, and 0.902 at *P* = 0.50. The full run comprised 1,800 successful model fits.

#### F.4 Selection rules and computation

In every experiment, the generating effect vector has exact zero sum and exact zeros outside the active set. Thus ***P β***^true^ = ***β***^true^, and the nonzero generating effects and nonzero centered contrasts define the same true active set. All selection metrics evaluate recovery of this set. PGL and PIND use the same likelihood, support information, scale priors, and selection rule; their paired comparison isolates the contribution of phylogenetic borrowing. Feature-wise comparisons evaluate signal recovery under each method’s normalization, zero handling, and testing procedure.

For selected set *Ŝ* and true support *S*, power is | *Ŝ ∩ S*|*/*|*S*|, FPR is | *Ŝ ∩ S*^*c*^|*/*|*S*^*c*^|, and the false discovery proportion (FDP) is | *Ŝ ∩ S*^*c*^|*/*| *Ŝ* |, defined as zero for an empty selection. FDR is estimated by the mean FDP over replicates. Metrics in the tables are averages of replicate-level metrics. The *F*_1_ score summarizes precision and recall. RMSE compares coefficient estimates with the identifiable, zero-sum true coefficient vector.

All Bayesian runs used 40,000 MCMC iterations, discarded the first 20,000 as burn-in, and retained every fourth remaining draw, leaving 5,000 draws. The global and local log-Cauchy priors and nuisance priors were those in Section 2 of the main article. Proposal scales were adapted during warm-up toward acceptance probability 0.40. We checked support, zero-sum and RMS invariants, completion and model errors, and sampler acceptance rates. For Experiments 2 and 3, recorded diagnostics also include coefficient effective sample size (ESS), Monte Carlo standard error (MCSE), and autocorrelation. Across their settings and methods, the means of the replicate-level median coefficient ESS ranged from 478 to 843 in Experiment 2 and from 232 to 989 in Experiment 3; the corresponding median coefficient MCSE means ranged from 0.0008 to 0.0137 and from 0.0029 to 0.0097. The recorded diagnostics for Experiment 1 include acceptance summaries but not coefficient ESS or MCSE summaries. The simulation results contain 1,400 replicates across all settings for Experiment 1, 800 for Experiment 2, and 1,800 model fits for Experiment 3, with no reported fit failures. Coefficient-level Monte Carlo precision, rather than mixing of the global scale alone, is the relevant diagnostic for the feature-specific comparisons.

### G Additional Real-Data Results

#### G.1 Cohort, preprocessing, and inferential target

The salivary 16S rRNA data were obtained from Jayaraman et al. (2025). The supplied phyloseq object contained 76 samples and 235 ASVs. After removing positive and negative control samples, we retained 26 participants living without HIV (group A) and 40 living with HIV (group B). We required detection in at least 10% of samples in each group, corresponding to at least 3 group-A and 4 group-B samples; 70 ASVs remained. This groupwise filter excludes ASVs never observed in one group, for which an observed-support abundance contrast would be especially weakly identified. It does not model prevalence or establish biological presence. The analysis used the supplied phylogeny and its Brownian correlation kernel. The clinician-assigned age groups were based on a 60-year cutoff; an indicator for the older group (coded 2) was included as a nuisance covariate. HIV status was coded as one for group B and zero for group A. A positive group coefficient indicates a larger log-ratio with HIV relative to the mean across retained ASVs, conditional on each sample’s observed support.

PGL and PIND used the same observed-support multinomial likelihood, zero-sum group effects, nuisance design, and shrinkage priors. The only difference was that PIND fixed *ω* = 0. We ran four independent chains per model for 80,000 iterations each, discarded the first 40,000 iterations, and retained every eighth subsequent draw. This gave 5,000 post-warm-up draws per chain and 20,000 combined draws per method. Selection required exclusion of zero by the 95% equal-tailed credible interval. DESeq2 and LinDA were fitted to the same samples and ASVs; their selections used Benjamini–Hochberg adjusted *p <* 0.05. All reported coefficients are expressed in natural-log units for the change from living without HIV to living with HIV. DESeq2 uses size-factor normalization, LinDA uses bias-corrected log-ratio regression with zero adjustment, and PGL and PIND use zero-sum contrasts conditional on observed support. These procedures differ in normalization, reference, zero treatment, and selection rules, so the cross-method comparisons below are descriptive.

#### G.2 Selection patterns and borrowing diagnostics

PGL selected 12 ASVs, PIND 9, DESeq2 8, and LinDA 18. All nine PIND selections were included among the PGL selections. Of the PGL selections, four overlapped DESeq2 and six overlapped LinDA; four ASVs were selected by both feature-wise procedures. The PGL and PIND posterior means had Pearson correlation 0.989 across all 70 ASVs, with identical estimated signs for 66 of them. The three PGL-only selections were ASVs assigned to *Oribacterium, Streptococcus*, and *Scardovia*; they had the same estimated direction under PIND, but their PIND intervals crossed zero. The different LinDA and DESeq2 selection sets should not be interpreted as a common FDR-controlled comparison with the Bayesian interval rule.

The posterior mean of PGL’s borrowing weight was 0.724 (median 0.782; 95% credible interval 0.166–0.991). The posterior probability that *ω >* 0.5 was 0.830. This broad distribution is compatible with partial borrowing and does not support a claim that the tree fully determines the group effects. The radial tree display in Figure S1 locates PGL selections but does not demonstrate phylogenetic clustering: descriptive lower-tail permutation probabilities for the mean pairwise distance of selected ASVs with positive and negative coefficient estimates were 0.739 and 0.457, respectively. We therefore do not use visual proximity of selected tips as independent validation of borrowing.

Rank-normalized split 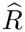 for every group coefficient was below 1.05 in both fits (maxima 1.023 for PGL and 1.040 for PIND). Median values were 1.004 in both fits, and the minimum between-chain correlation of the 70 coefficient means exceeded 0.99 for each method. The borrowing-weight split 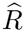 was 1.004. These checks support the reported group-effect summaries; they do not establish biological replicability in another cohort.

#### G.3 The *Leptotrichia* ASV comparison

For the *Leptotrichia* ASV discussed in the main text, PGL and PIND posterior means were −0.952 and −0.950, with 95% credible intervals [−1.467, −0.423] and [−1.459, −0.411], respectively. LinDA estimated 1.896 (BH-adjusted *p* = 0.00205), and DESeq2 estimated 0.458 (BH-adjusted *p* = 0.532). The detection counts and mean relative abundances among samples in which the ASV was detected are unadjusted descriptive summaries in the main text; relative abundance uses each sample’s total count across the 70 retained ASVs as its denominator. These summaries are distinct from the age-adjusted log-ratio coefficients. The sign difference accompanies changes in zero treatment and compositional reference; this comparison does not separate their individual contributions.

**Figure S1:**
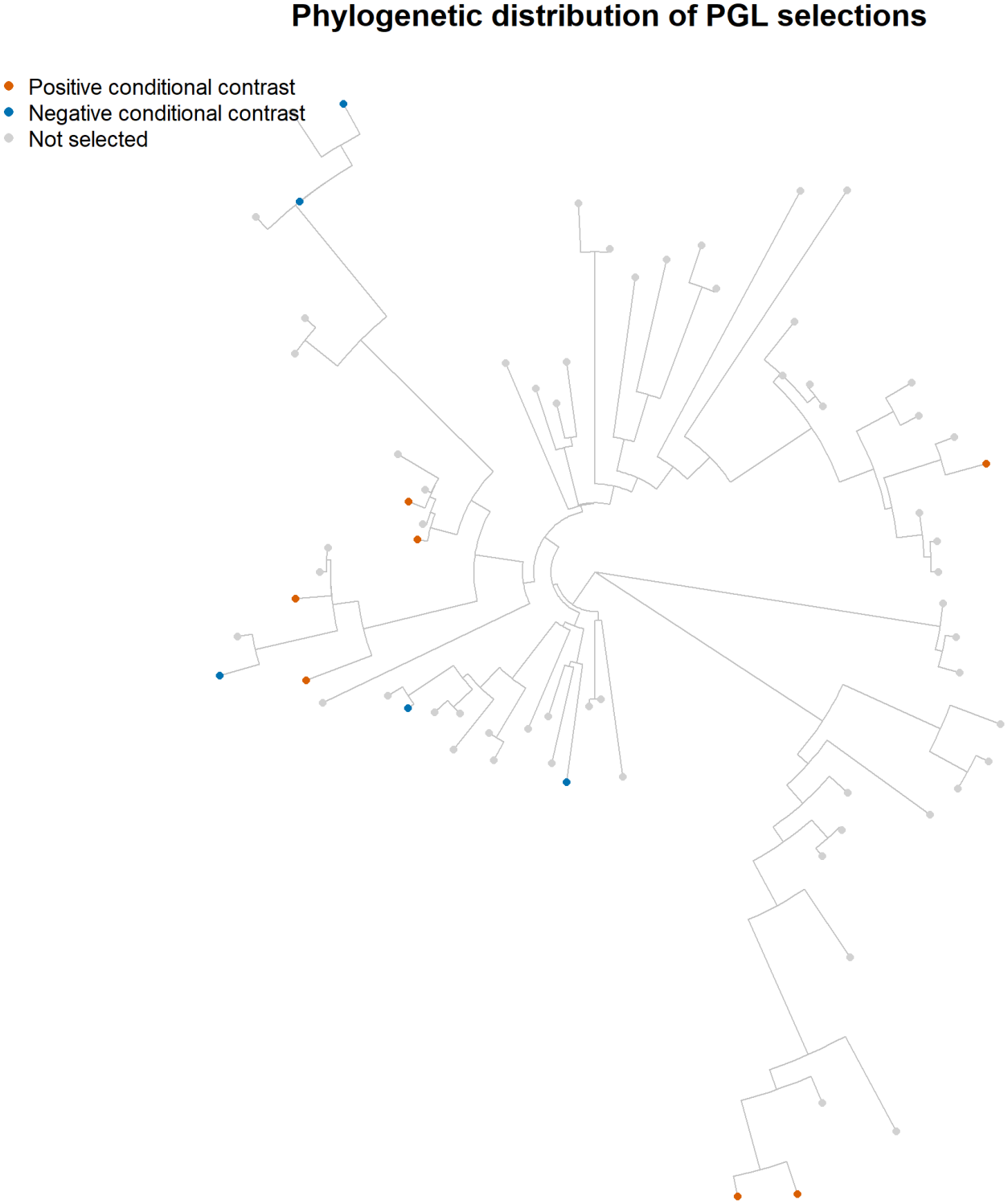
Descriptive distribution of the 12 PGL selections on the phylogenetic tree of microbial ASVs used in the HIV oral microbiome analysis. Orange and blue tips have positive and negative regression coefficient estimates for HIV status, respectively; gray tips were not selected. The display does not show statistically significant clustering of selected tips (permutation results in the text).

#### G.4 Complete ASV-level summaries

Table S8 reports the PGL and PIND posterior means, 95% credible intervals, and selection status for every retained ASV. Table S9 reports the corresponding DESeq2 and LinDA estimates in natural-log units and adjusted *p* values. Each table row represents one ASV, labeled by its genus or the lowest available taxonomic rank; repeated names denote distinct ASVs, without taxonomic aggregation. Rows appear in the same order in both tables. The replication files link table rows and figure positions to the original feature identifiers and taxonomy. The machine-readable results also include posterior medians, false-sign probabilities, and coefficient-level split 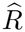.

**Table S8:**
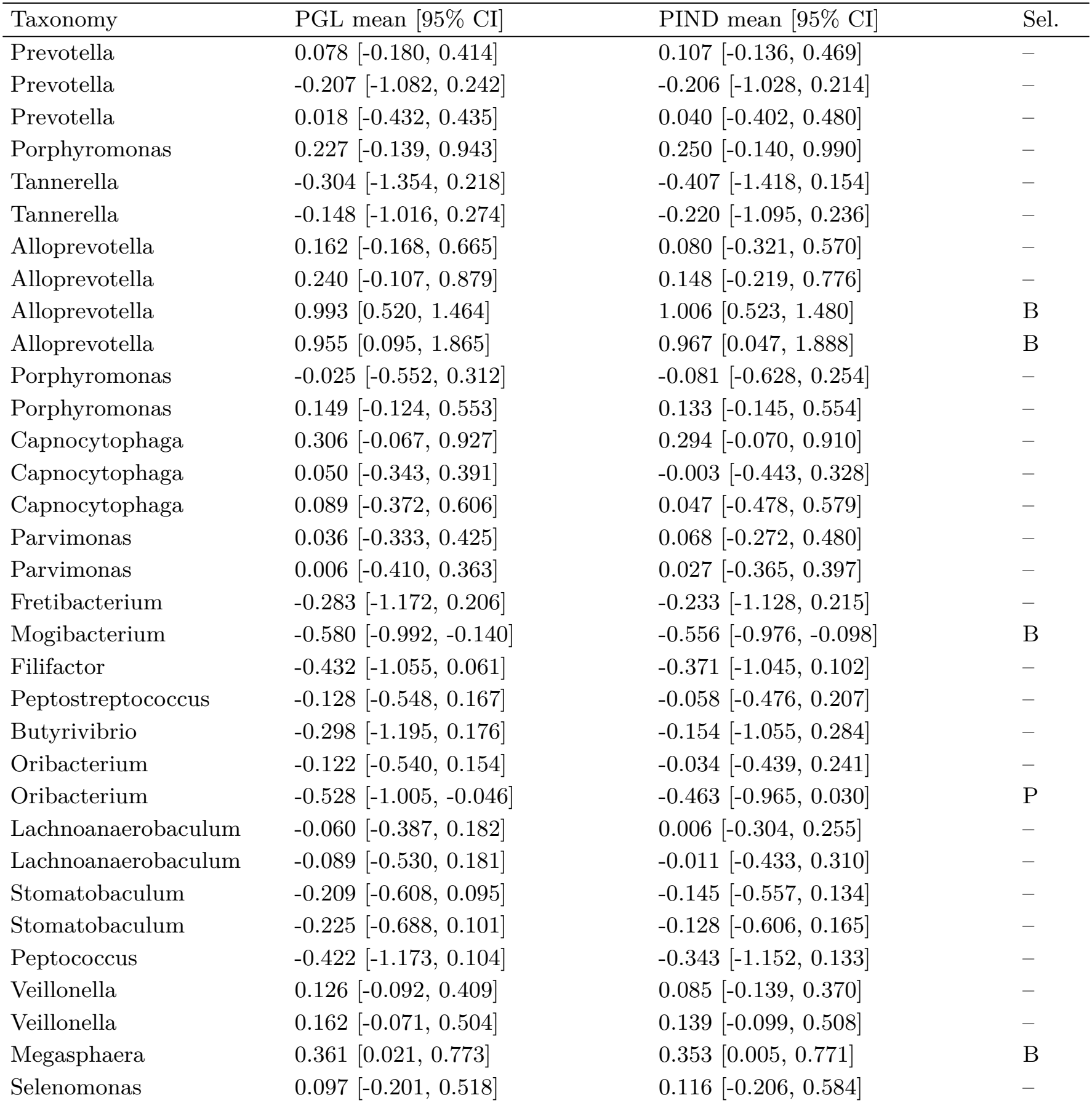
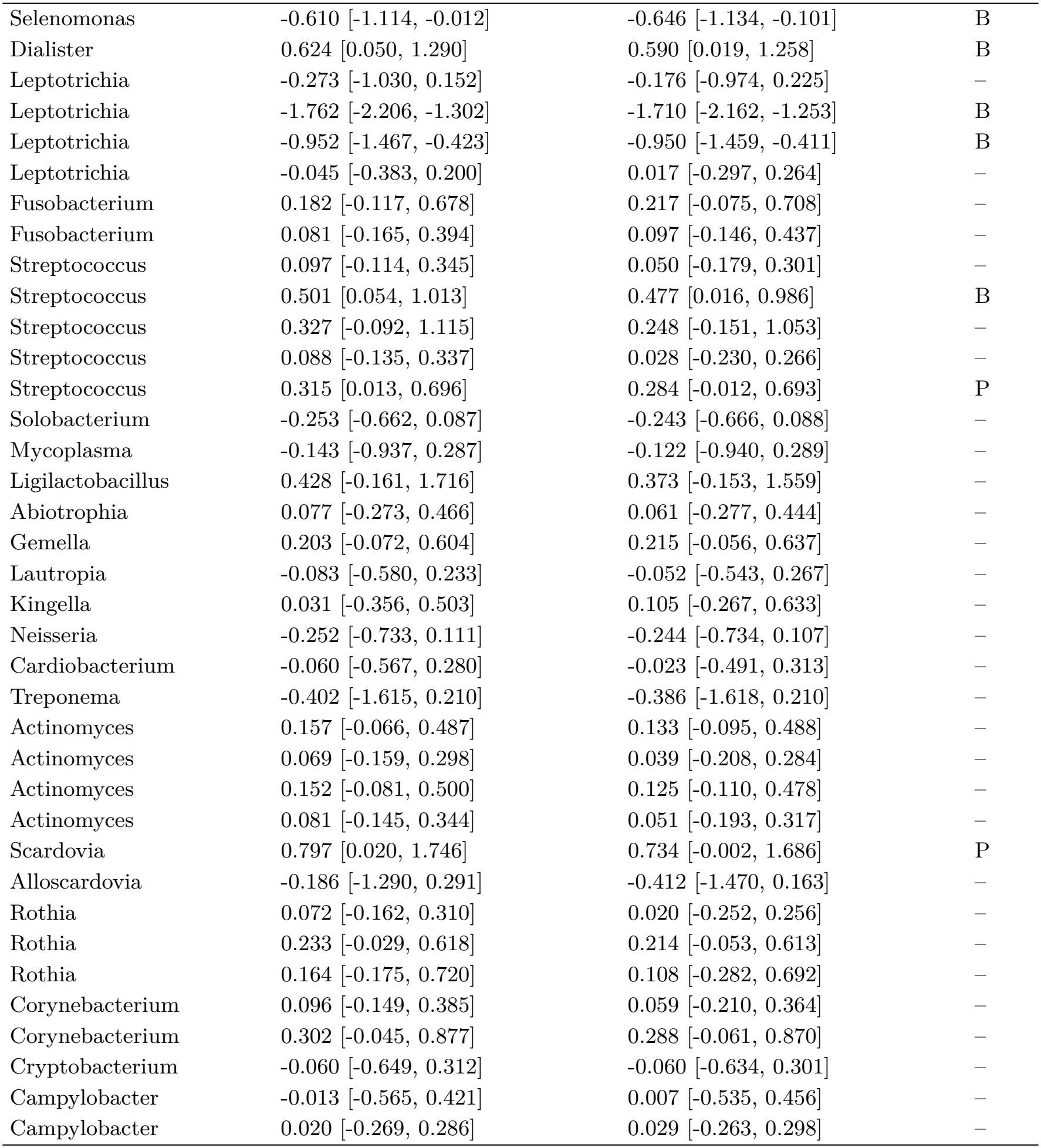
Complete Bayesian summaries of the regression coefficients for HIV status for all ASVs retained in the HIV oral microbiome analysis. Posterior means and 95% equal-tailed credible intervals are shown; B denotes selection by both PGL and PIND, P denotes PGL only, and a dash denotes neither. Effects are zero-sum log-ratio contrasts for HIV status, conditional on observed support. Each row represents one ASV; repeated taxonomic names denote distinct ASVs.

**Table S9:**
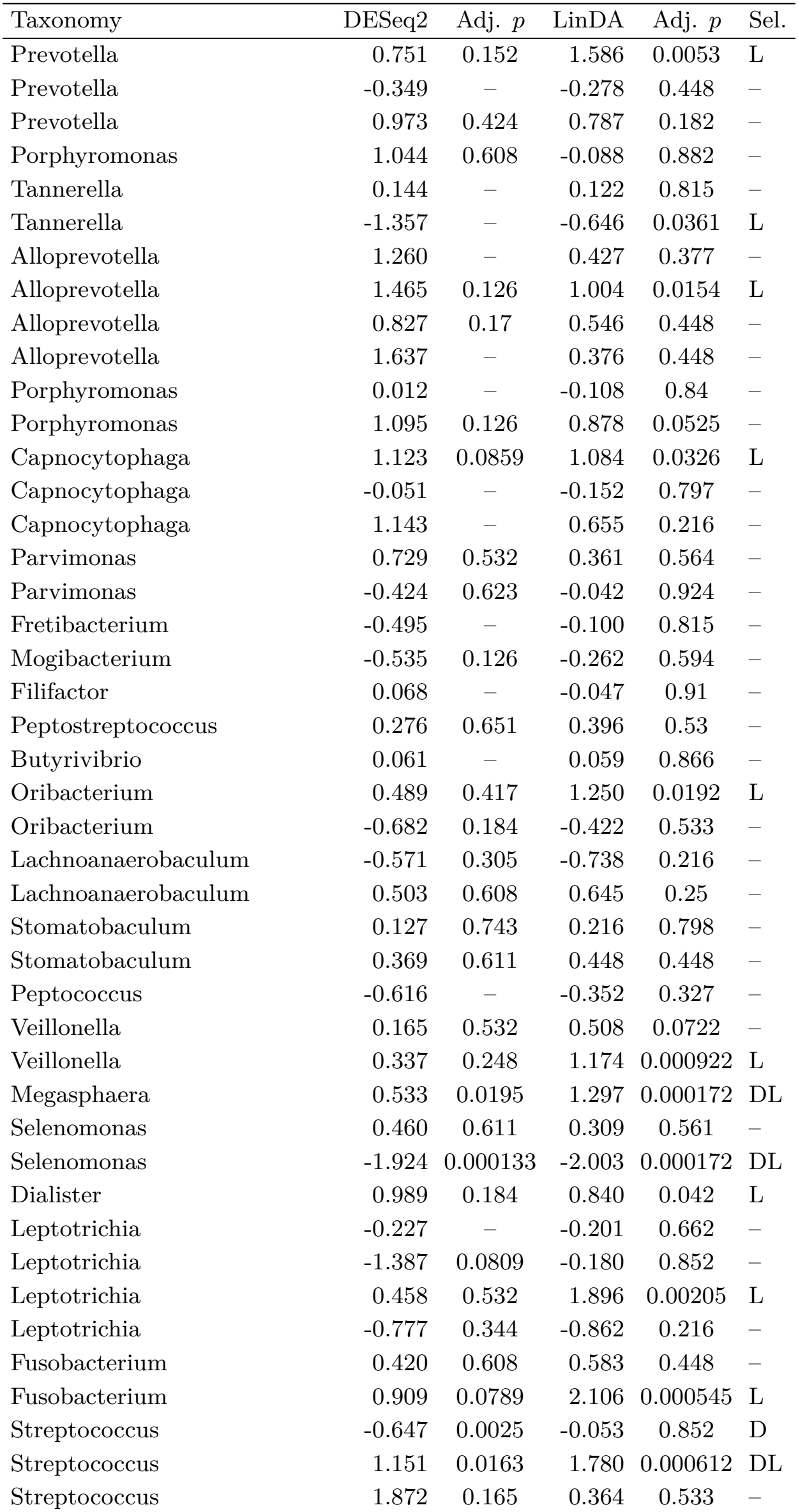
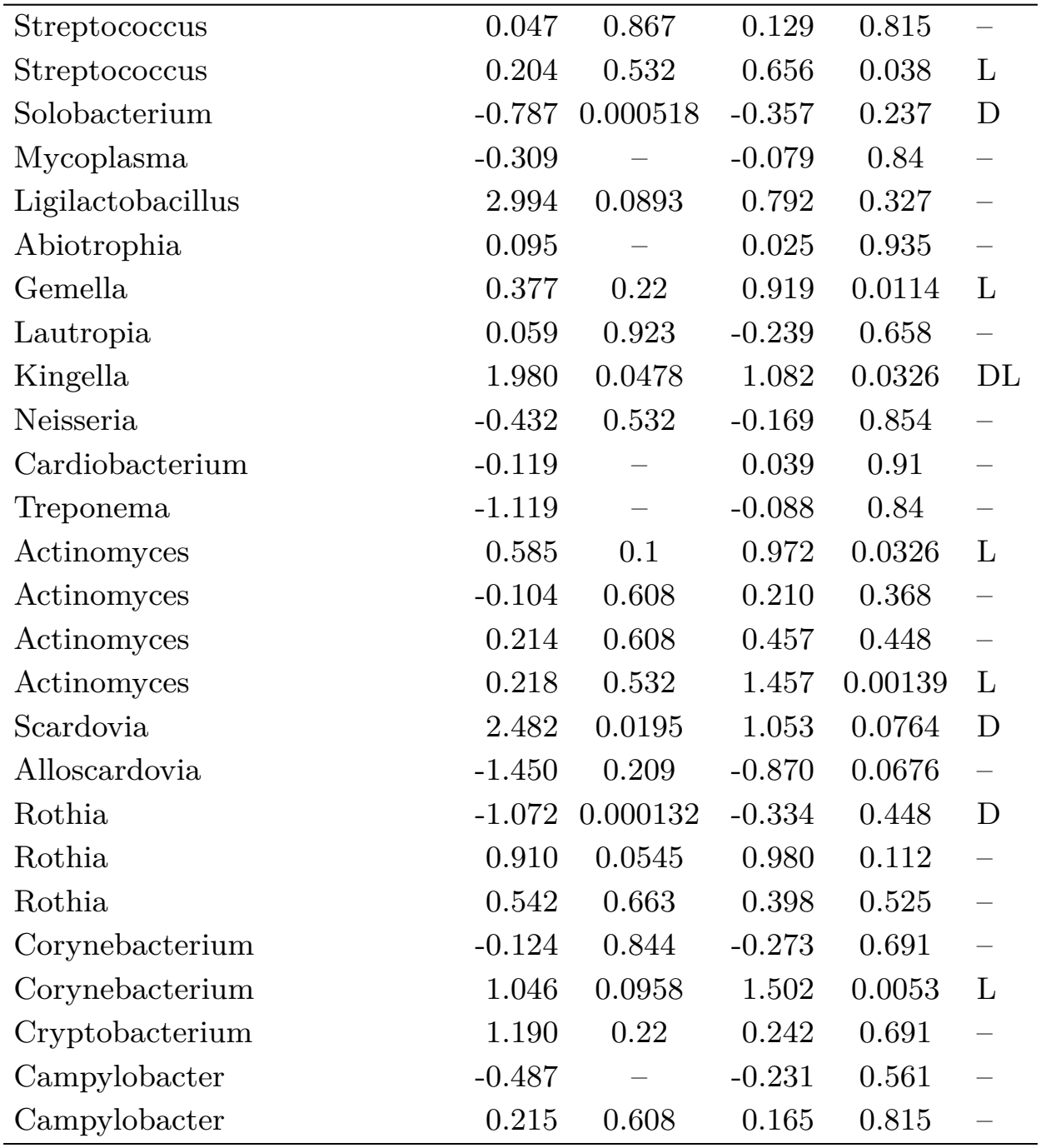
Complete feature-wise benchmark summaries for the same 70 retained ASVs. Estimates and BH-adjusted *p* values are shown. A dash in an adjusted p-value column indicates an unavailable value; a dash in the selection column indicates selection by neither method. D and L in the last column mark DESeq2 and LinDA selections at adjusted *p <* 0.05, respectively. Estimates are in natural-log units for HIV status; normalization, reference, and zero treatment differ across methods. Each row represents one ASV, in the same order as the Bayesian table; repeated taxonomic names denote distinct ASVs.

